# X-CRISP: Domain-Adaptable and Interpretable CRISPR Repair Outcome Prediction

**DOI:** 10.1101/2025.02.06.636858

**Authors:** Colm Seale, Joana P. Gonçalves

## Abstract

**Motivation:** Controlling the outcomes of CRISPR editing is crucial for the success of gene therapy. Since donor template-based editing is often inefficient, alternative strategies have emerged that leverage mutagenic end-joining repair instead. Existing machine learning models can accurately predict end-joining repair outcomes, however: generalisability beyond the specific cell line used for training remains a challenge, and interpretability is typically limited by suboptimal feature representation and model architecture.

**Results:** We propose X-CRISP, a flexible and interpretable neural network for predicting repair outcome frequencies based on a minimal set of outcome and sequence features, including microhomologies (MH). Outperforming prior models on detailed and aggregate outcome predictions, X-CRISP prioritised MH location over MH sequence properties such as GC content for deletion outcomes. Through transfer learning, we adapted X-CRISP pre-trained on wild-type mESC data to target human cell lines K562, HAP1, U2OS, and mESC lines with altered DNA repair function. Adapted X-CRISP models improved over direct training on target data from as few as 50 samples, suggesting that this strategy could be leveraged to build models for new domains using a fraction of the data required to train models from scratch.

**Availability:** An implementation of X-CRISP is available at github.com/joanagoncalveslab/xcrisp.

## Introduction

Gene therapies that alter the DNA to treat diseases have been made widely accessible with the emergence of CRISPR (Clustered Regularly Interspaced Short Palindromic Repeats) technology (Jinek *et al*., 2012), providing faster, cheaper, and more effective gene editing (Bhattacharya *et al*., 2015; Maxmen, 2019; Wang *et al*., 2014; Hsu et al., 2014; Adli, 2018). The CRISPR strategy to gene editing performs enzyme-based cleavage of the DNA at a programmed location, determined by a guide RNA sequence, and subsequently exploits endogenous mechanisms recruited by the cell to repair the DNA break and introduce the desired changes. The success of gene therapies relies on CRISPR editing to produce a precise outcome, regardless of whether aiming to inactivate a disease-causing gene, to replace such a gene with a healthy copy, or to introduce a new gene with therapeutic properties. In principle, homology-directed repair (HDR) offers the most control over the repair outcome, given that it can make use of a donor template. However, HDR is typically inefficient, as it is only available during the G2 and S phases. As an alternative, template-free editing can be performed throughout the cell cycle, leading to repair by HDR or one of the more error-prone non-homologous end joining (NHEJ) and microhomology-mediated end joining (MMEJ) pathways (Scully *et al*., 2019).

Template-free editing is appealing for its wide availability but presents challenges to ensure a precise post-repair outcome, given the stochasticity of the repair processes as a result of pathway choice and inaccuracies of the repair machinery. Notably, numerous studies have reported a strong dependence of CRISPR-induced repair outcomes on the DNA sequence surrounding the break site (Koike-Yusa *et al*., 2014; van Overbeek *et al*., 2016). This suggests that it might be possible to influence the post-repair outcome distribution by purposefully designing guide RNAs to target sequence contexts favouring desired outcomes.

Several machine learning models leverage the relationship with sequence context to predict the frequency distribution of CRISPR-induced DNA repair outcomes, aimed at improving guide RNA design. We categorise these models regarding predicted outcomes into more or less granular. The more granular models estimate the distribution of individual repair outcomes, and include: inDelphi, using a dual neural network and k-nearest neighbours model (Shen *et al*., 2018); and FORECasT (Allen *et al*., 2019) and Lindel (Chen *et al*., 2019), both multinomial logistic regression models. The less granular models predict the frequencies of aggregated or higher-level repair outcomes: SPROUT, using a gradient-boosted tree (Leenay *et al*., 2019); and CROTON, based on a convolutional neural network (Li *et al*., 2021). Less granular models lack detail to offer control over precise repair outcomes. For example, CROTON predicts the overall frequency of 1bp insertions, whereas inDelphi predicts a 1bp insertion frequency per nucleotide.

Model interpretability is another key aspect of repair outcome prediction that has been insufficiently explored. The ability to explain predictions for individual target sequences and delineate how features such as sequence properties influence changes in outcome frequency provides a means to scrutinise model output and gain insight into DNA repair processes, as well as to optimise gene editing. However, less granular models make it infeasible to explain individual outcomes, whereas more granular models show a tradeoff between interpretability and performance. Specifically, FORECasT and Lindel outperform inDelphi (Chen *et al*., 2019), but are difficult to interpret due to the use of a large number of features with suboptimal encoding. Despite their linear model architecture, FORECasT and Lindel rely on over 3000 binary features related to sequence context and repair outcome characteristics. Furthermore, features such as deletion or microhomology length are one-hot encoded, making it challenging to recover the inherent relationship between different values of the same feature. This also leads to significant sparsity, with many features showing marginal contributions to a large proportion of outcomes. The inDelphi model uses non-linearity to leverage a more compact feature set, but ignores microhomology location and increases model complexity by distributing the prediction of deletion outcomes across multiple models. We argue that improved interpretability could be achieved by pairing a compact set of interpretable features with a non-linear model, while keeping model complexity as low as possible.

Repair outcomes are further influenced by cellular and genomic context.However, training models for each context requires considerably large data that might be unavailable or challenging to generate. Notably, most pre-trained models have been trained on a few mice and human cell lines, for which data either previously existed or was purposely generated (Shen *et al*., 2018; Allen et al., 2019; Chen et al., 2019; Leenay et al., 2019; Li et al., 2021). The limited diversity of genomic contexts covered by current prediction models can affect the reliability of model prediction in new genomic contexts. Combined with the high cost of generating new data, this could impede model adoption in more diverse or unique genomic contexts, such as those encountered when developing CRISPR therapeutics (Chehelgerdi *et al*., 2024) for precision medicine or rare diseases. To address this challenge, we explore transfer learning (TL) as a means to reuse and adapt knowledge from repair outcome prediction in data-rich genomic contexts for prediction in new contexts with limited data availability (Pan and Yang, 2009; Weiss *et al*., 2016; Tan et al., 2018). The success of TL is influenced by the degree of “relatedness” between the source and target prediction domains, while evidence suggests that DNA repair mechanisms are highly conserved among eukaryotes (Lehmann and Taylor, 2001; Cahill et al., 2006) and that models trained on mouse embryonic stem cell (mESC) data can reasonably predict outcomes in zebrafish and *Xenopus* embryos (Naert *et al*., 2020). We hypothesise that TL could exploit the conservation of DNA repair mechanisms to facilitate adaptation of pre-trained CRISPR outcome prediction models to data-scarce genomic contexts.

Here, we introduce X-CRISP (eXplainable CRISPR Predictions), a repair outcome prediction model designed to be granular, interpretable, and sufficiently flexible to enable adaptation to new genomic contexts. X-CRISP integrates a neural network model based on five deletion-descriptive features for prediction of deletion outcomes, alongside two multinomial logistic regression models for prediction of insertion outcomes and deletion-insertion ratio. We employ Shapley values (Lundberg and Lee, 2017) to interpret the behaviour of X-CRISP for each outcome prediction. Finally, we demonstrate several transfer learning strategies, wherein X-CRISP models pre-trained on wild-type (WT) mESC cells are adapted to other domains encompassing different cell types, organisms, and genotypes with altered DNA repair function.

## Methods

### Data and preprocessing

We used sequence data from two template-free CRISPR targeting screens: FORECasT ([dataset]* Allen *et al*., 2018) and inDelphi ([dataset]* Shen *et al*., 2018). Both studies employed thousands of designed gRNAs paired with a 55bp (inDelphi) or 79bp (FORECasT) DNA sequence containing a PAM-adjacent 20bp target, which were delivered to Cas9-expressing cells via lentiviral transduction. Following several days of cell culture for genomic integration, DNA cleavage, and repair, DNA sequencing was performed to capture the CRISPR repair products.

#### Targets, samples, and outcome sequence data

From FORECasT, we analyzed the 11,058 “Explorative gRNA-Targets” in the “FORWARD” orientation (“NGG” PAM, not “CCN”), requiring a minimum of 30bp on both sides of the cut site. From inDelphi, we used the 1,996 “FORWARD” gRNA-target pairs in “Lib-A”. Specifically, we examined data from mouse embryonic stem cells (mESCs), either in their wild-type form (WT) or upon double-knockout of Prkdc and Lig4 (denoted by *Prkdc*^*-/-*^*Lig4*^*-/-*^) resulting in NHEJ deficiency (hereafter denoted by *NHEJ*^*-/-*^, Yue *et al*. (2020)). We also included data from human leukemic near-haploid cells (HAP1), human osteosarcoma cells (U2OS), and modified human chronic myelogenous leukaemia cells (TREX2), where the TREX2 modification fuses the Cas9 protein to the three-prime repair exonuclease 2. All FASTQ files were obtained from the European Nucleotide Archive (Leinonen *et al*., 2010) (see Supplementary Table S1 for study and accession numbers).

#### Sequence alignment, repair outcome calling

We relied on the same set of tools to process all datasets. For FORECasT data, we used PEAR v0.9.11 (Zhang *et al*., 2014) to merge paired-end reads using parameters “-n 20 -p 0.1” (specifying a minimum combined sequence length of 20 and a probability of no overlap below 0.1) and the “indelmap” tool (Allen *et al*., 2019) to map the merged reads to target sequences, both parameterised and performed as in Allen *et al*. (2019), also discarding reads mapped to multiple targets. For inDelphi, we reverse complemented the target-containing reverse reads before mapping, also using the same “indelmap” tool as described in Allen *et al*. (2019) (Supplementary Fig. S1-S6 show the distributions of counts of reads mapped to target sites per dataset). Finally, we used SIQ v4.3 (van Schendel *et al*., 2022) to call repair outcomes per read with options “-m 2 -c -e 0.05”, specifying a minimum number of 2 reads for the event to be counted, the collapsing of identical events to a single record with the corresponding sum of counts, and a maximum permitted base error rate of 0.05.

#### Repair outcome profile generation

We calculated outcome frequency distributions (or repair outcome profiles) per gRNA-target and screen as follows. We considered all deletions up to 30bp long overlapping the cut site or adjacent to the upstream nucleotide neighbouring the cut site. This cut-off was selected because deletion lengths over 30 bp were rarely observed (Supplementary Fig. S7–S12), and also to maintain consistency with other studies on CRISPR repair outcome profile prediction. (Allen *et al*., 2019; Chen *et al*., 2019). We then determined unique deletion outcomes by grouping deletions that produced identical repair products. Only MH-*based* deletions presented such ambiguity, as a result of the loss of one of two microhomologous sequences flanking the deleted region (Fig. 1). All remaining unique deletions were categorised as “MH-*less* deletions”. This yielded unique sets of approximately 330-480 deletion outcomes per target site. We also considered all unique single- and di-nucleotide insertions, as well as one single category for insertions of at least 3bp due to their rarity of occurrence (Supplementary Fig. S13-S18), totalling 21 insertion outcomes. We mapped reads to outcomes per gRNA-target, and discarded gRNA-targets with less than 100 mutated reads. Outcome counts were divided by the sum of counts per gRNA-target to obtain the final outcome profiles. The original target sequences (and their respective repair outcome profiles) were randomly split into non-overlapping train and test sets per source study, FORECasT and inDelphi, to guarantee that train and test sets were disjoint for all datasets of that study. The inDelphi WT mESC dataset itself was not split, given that it was only used to test models trained on the FORECasT WT mESC train set. For the four datasets used as contexts for adaptation with transfer learning (i.e. U2OS, HAP1, *NHEJ*^*-/-*^, TREX2), we obtained smaller train sets of 500 sequences by random subsampling the larger train set following the split per study. Train sets were later used for model hyperparameter optimisation and test sets were held out for evaluation (Table 1). The small variations in the numbers of test sequences across datasets from the same study in Table 1 result from the filtering. We calculated Needleman-Wunsch (Needleman and Wunsch, 1970) and Smith-Waterman (Smith *et al*., 1981) alignment scores (scoring: match=1.0, mismatch=0.0, gap=0.0), and Hamming distances between sequences in the train and test sets to ensure there were no near-identical sequences between them (Supplementary Fig. S19-S21).

**Fig. 1:**
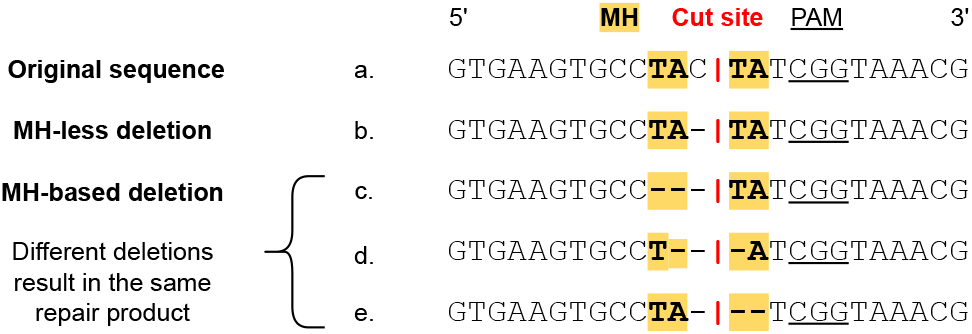
Deletion type categorisation.

**Table 1.** Post-filtering counts of processed repair outcome profile datasets per cell line, genotype, and study. Total counts, as well as train and test set splits.

| Cell line | Genotype | Study | Total | Train | Test |
| --- | --- | --- | --- | --- | --- |
| mESC | WT | FORECasT | 9854 | 5900 | 3954 |
| mESC | WT | inDelphi | 1961 | 0 | 1961 |
| mESC | *NHEJ*<sup>-/-</sup> | inDelphi | 1485 | 500 | 985 |
| K562 | TREX2 | FORECasT | 3855 | 500 | 3355 |
| HAP1 | WT | FORECasT | 4450 | 500 | 3950 |
| U2OS | WT | inDelphi | 1462 | 500 | 962 |

**Table 2.** X-CRISP deletion outcome feature descriptions.

| Feature | Description |
| --- | --- |
| MH length | Length of MH, defined as $M \in [0, 30]$. Zero for MH-less deletions. For MH-based deletions, $M \in [1, 30]$, since the deletion contains one of the MHs (Fig. 1) and 30 is the maximum deletion length. |
| MH GC | Fraction of GC in MH, defined as $F \in [0, 1]$. Zero for MH-less deletions. |
| Gap | Length between MHs, defined as $G \in [1, 30]$. For MH-less deletions, it equals the deletion length. |
| Left edge | Deletion left edge position $L \in [-30, 0]$, where 0 indicates the edge is at the cut site. For MH-based deletions: $L$ is the distance in bp from the cut site to the closest base pair of the PAM-distal MH. |
| Right edge | Deletion right edge position $R \in [0, 30]$, where 0 indicates the edge is at the cut site. For MH-based deletions: $R$ is the distance in bp from the cut site to the closest base pair of the PAM-proximal MH. |

### X-CRISP

The proposed model, X-CRISP, uses three sub-models to predict the repair outcome profile for a 60bp sequence centred at the cut site. The first two sub-models predict individual deletion and insertion outcomes, and the third predicts the overall frequency of deletions and insertions. The outputs of the first two sub-models are scaled by the output of the third and concatenated to construct the complete predicted repair outcome profile.

#### Deletion model

The deletion model predicts a frequency distribution per target over all considered deletion outcomes. Different from the other granular deletion models, X-CRISP introduces common features for MH-*based* and MH-*less* deletions, and avoids one-hot encoding by representing integer features as-is. Five features are reconciled and used ubiquitously by X-CRISP across deletion categories to consolidate deletion prediction into one single interpretable model: “Left edge” and “Right edge”, representing the left and right deletion edges or the positions of the nucleotides closest to the cut site for the left and right MHs; “Gap”, denoting the distance between the two edges; and MH length and MH GC fraction, which are both zero for MH-*less* deletions. These features are fed to a fully connected neural network that independently scores each outcome between 0 and 1. The network contains two hidden layers of 16 nodes and one output node, using sigmoid activation at every layer. We trained two models, “X-CRISP KLD” and “X-CRISP MSE”, respectively using the Kullback-Leibler divergence (KLD) (Kullback and Leibler, 1951) and mean-squared error (MSE) loss functions. Training was performed using PyTorch v.1.8.0 with the Adam optimiser (Kingma and Ba, 2014) (*β*_1_ = 0.99, *β*_2_ = 0.999, and remaining default settings), a batch size of 200, and learning rates 0.05 and 0.01 for KLD and MSE, respectively. We applied an exponential learning rate decay with *γ* = 0.999 per epoch. We used L2 regularisation and optimised hyperparameters with 5-fold cross-validation (CV) on the train set. The final models were trained on the entire train set using the hyperparameter values yielding the lowest mean loss (Supplementary Table S2 for tested and final hyperparameters).

#### Insertion and deletion-insertion models

The insertion model predicts frequencies for the 21 insertion outcomes, while the deletion-insertion model predicts the overall frequency of deletions and insertions. Both models use softmax regression and take a DNA sequence as input, represented by one-hot encodings of single nucleotides and dinucleotides at each position. The insertion model uses the six nucleotides directly upstream of the PAM, and the deletion-insertion model considers the 20bp target sequence. Both models were trained using the Adam optimiser and an exponential learning rate decay. We used L2 regularisation and optimised hyperparameters using 5-fold CV on the train set. The final models were trained on the entire train set using the hyperparameter values yielding the lowest mean MSE (Supplementary Table S2 for tested and final hyperparameters). We trained for a maximum of 200 epochs, with early stopping if there was no improvement after two epochs.

#### Other prediction models

We compared X-CRISP against four other published models at the time of writing: inDelphi (Shen *et al*., 2018), FORECasT (Allen *et al*., 2019), Lindel (Chen *et al*., 2019), and CROTON (Li *et al*., 2021) (Supplementary Table S3). We excluded SPROUT (Leenay *et al*., 2019), as it only predicts aggregate outcomes at a higher level, namely: “average insertion length”, “average deletion length”, “diversity”, and “most likely inserted pair”. These prediction tasks do not align well with the practical applications we envision for X-CRISP, which require more detailed outcomes. Every model was trained on the same data (Table 1), following the procedures outlined in the respective publication, with two exceptions: we trained inDelphi with a maximum deletion length of 30bp for consistency across models, and we trained CROTON using the architecture provided by the authors without redoing the architecture search.

#### Evaluation of prediction performance

We trained each model on 5900 FORECasT WT mESC target repair profiles and tested it against 3954 FORECasT and 1961 inDelphi target repair profiles, without overlap between train and test. Direct comparisons were challenging due to the different outcome categorisations used by each model (Supplementary Table S3). To address this, we assessed each model on the outcomes described in its original publication, as well as on three sets of outcomes comparable across models: common MH-*based* deletions, common MH-*less* deletions, and 1bp insertions. Across models, predicted deletion outcomes were limited to 30bp in size. For comparison with FORECasT for 1bp insertions, non-repeat insertions of nucleotides neighbouring the cut site were grouped as one outcome. To measure the error between predicted and observed repair outcome probability distributions, we used the Jensen-Shannon distance (JSD) and Pearson’s correlation coefficient. The JSD is designed to measure the divergence between probability distributions, and its symmetry and bounded range (0 to 1) make it respectively invariant to the role of the two distributions and more comparable across targets with varying numbers of possible repair outcomes than other relevant metrics such as KLD or cross-entropy. The Pearson’s correlation quantifies the linear relationship between the predicted and observed frequency vectors of each target. It is less suitable for probability distributions because it assumes independence between the values in each vector, whereas the probabilities associated with the repair outcomes of any target sum to 1 and are necessarily dependent on one another. Nevertheless, Pearson’s correlation is still commonly used and we include it for completeness. For the overall frequencies of deletions and insertions, we used MSE.

We also assessed the ability of each model to classify “precision-X%” target sequences, defined as those where a single outcome represented at least *X*% of the reads assigned to all considered repair outcomes, with *X* ∈ {20, 30, 40, 50, 60, 70}. Targets were classified as positive if they met the criteria and negative otherwise. We used precision, recall, and Matthew’s correlation coefficient (MCC) to evaluate performance. Precision denotes the rate at which the model is expected to be correct when it predicts a target sequence as a precision-X% site. Recall expresses the fraction of all observed precision-X% targets of a set we can expect the model to identify. For CRISPR-based gene editing, the focus is on which targets can be more confidently used to produce the desired outcome with the highest possible fidelity for more precise gene editing. As a result, higher emphasis is placed on precision and reducing the risk of false precision-X% target predictions, compared to recall and recovering the most precision-X% targets. The MCC evaluates the quality of the binary precision-X% predictions made by the model, including but also beyond precision and recall, providing a value between −1 and +1, where +1 indicates perfect agreement, 0 indicates random prediction, and −1 indicates total disagreement between model predictions and observed outcomes. The MCC is generally reported as more informative and reliable to assess binary classification than other combined performance metrics, such as F1 and AUC. It is also more suitable for evaluation under class imbalance, which is present in the case of precision-X% prediction tasks, where most targets are non-precision-X%. Lastly, we assessed six aggregate prediction tasks (deletion, 1bp insertion, 1bp deletion, 1bp frameshift, 2bp frameshift, and frameshift frequencies) using the MSE and Pearson’s correlation. We emphasise the MSE to determine the best performing model for these prediction tasks because MSE takes the magnitude of the errors into account, while Pearson’s correlation does not (it is both scale- and shift-invariant).

#### Explainability

To interpret X-CRISP predictions, we used SHapley Additive exPlanations (SHAP, python library v0.39.0 (Lundberg and Lee, 2017)). We calculated SHAP values to estimate the contribution of each feature to the change in outcome frequency predicted by the X-CRISP model for a given target, relative to the average frequency obtained for a background set of targets. For the deletion model, we used the “DeepExplainer” function with 10k randomly selected deletions from target sequences in the train set as background. For the insertion and deletion-insertion models, we used the “LinearExplainer” with 5k randomly selected targets as background. All other parameters were set as default. We generated SHAP values to explain 400 randomly selected target sequences from the test set per model. We aggregated the SHAP values for the nucleotide features by summing the SHAP values of their binary encodings. For example, the SHAP value for “A” at position 16 is the sum of the SHAP values for the feature|value pairs (A16|1, C16|0, G16|0, T16|0).

#### Transfer learning

We investigated whether X-CRISP models trained on FORECasT WT mESC data as a source domain could be adapted using transfer learning (TL) to predict on the following different cell lines as target domains: mESC *NHEJ*^*-/-*^, TREX2, HAP1, and U2OS. Our general TL approach was to first initialise a new model using the learned weights from the pre-trained X-CRISP KLD mESC model, and then either fine-tune (FT) or retrain and fine-tune (PF0-2) the initialised model using *n* samples from the other cell line of interest to adapt the model to the target domain. We used sets of samples of increasing size, with *n* ∈ {2, 5, 10, 20, 50, 200, 500}, where each subsequent set was a superset of the preceding one. Fine-tuning alone (FT) involved training the model on the target domain samples using a low learning rate. When retraining before fine-tuning (PF0-2), we controlled the flexibility of the model to adapt itself i.e. the number of learnable model parameters or weights) by freezing the weights of none (PF0), the first (PF1), or both (PF2) of its hidden layers before retraining, allowing only unfrozen layers to change (Tan *et al*., 2018) (Supplementary Table S2 for hyperparameter details). As baselines for comparison, we used the X-CRISP KLD model trained only on the WT mESC source domain data (SO) and another X-CRISP KLD model trained only on the target cell line data (TO) (Table 3).

**Table 3.** Baseline X-CRISP models and transfer learning strategies. Baselines: SO, source only; TO, target only. Transfer learning: FT, pretrained on source and fine-tuned for target; PF0-2, pre-trained on source and retrained + fine-tuned on target with 0-2 frozen hidden layers.

| Code | Weight initialisation | Frozen layers | Trained & fine-tuned on target |
| --- | --- | --- | --- |
| SO | Random | No | No |
| TO | Random | No | Both |
| FT | Pre-trained | No | Fine-tuned |
| PF0 | Pre-trained | No | Both |
| PF1 | Pre-trained | First hidden | Both |
| PF2 | Pre-trained | Both hidden | Both |

## Results and Discussion

### X-CRISP accurately predicts detailed repair profiles

We first evaluated the ability of all models trained on the 5900 FORECasT mESC target sequences to predict repair outcome profiles for the 3954 and 1961 mESC WT target sequences in the FORECasT and inDelphi test sets, respectively.

When predicting the profile as described in the original publication (Fig. 2, Original outcomes; Supplementary Fig. S22 for Pearson’s correlation), both X-CRISP KLD and X-CRISP MSE achieved the best performances with a significantly lower median JSD compared to FORECasT, the best non-X-CRISP model (FORECasT|inDelphi mESC data X-CRISP KLD: 0.43|0.37, FORECasT: 0.47|0.42; Wilcoxon two-sided signed rank test *p*-values *<* 0.05).

**Fig. 2:**
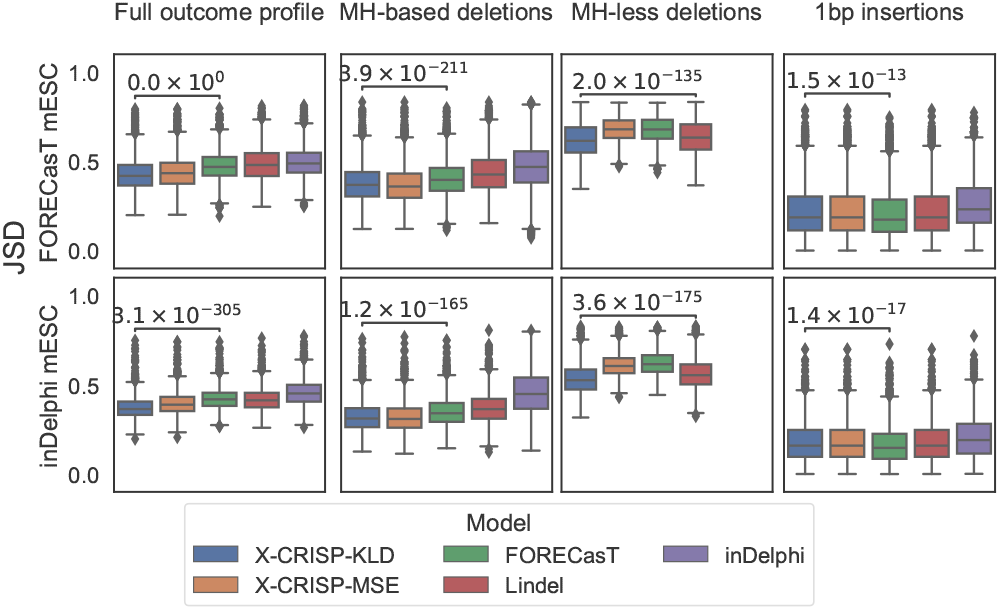
Detailed repair outcome prediction performance. Jensen-Shannon distance (JSD) between predicted and observed outcomes for (top) 3954 FORECasT or (bottom) 1961 inDelphi mESC WT test target sites, considering: (left to right) original publication outcomes; common MH-*based* deletions; common MH-*less* deletions; 1bp insertions. Significance *p*-values calculated using Wilcoxon two-sided signed-rank tests, comparing X-CRISP KLD to the best of the non-X-CRISP models. For 1bp insertions, X-CRISP KLD and Lindel perform identically, given that they are based on the same model, so the comparison is then made with FORECasT, the next best of the non-X-CRISP models.

For deletion frequency prediction, the X-CRISP KLD model significantly outperformed all others, with FORECasT and Lindel as the best non-X-CRISP models for MH-*based* (FORECasT|inDelphi mESC data X-CRISP KLD: 0.38|0.32, FORECasT: 0.40|0.35) and MH-*less* deletions (FORECasT|inDelphi mESC data X-CRISP KLD: 0.61|0.53, Lindel: 0.62|0.56), respectively (Fig. 2). The FORECasT and Lindel models could be at a disadvantage compared to X-CRISP due to their linear architectures and binary feature representations, since the one-hot encoding of integer-valued features (such as deletion length) and assumption of feature independence impairs their ability to model relationships between values within and across features. In contrast, X-CRISP uses integer-valued features as-is to safeguard interpretation, and leverages non-linearity to learn feature interactions. The inDelphi model uses a similar strategy, however X-CRISP additionally considers deletion edge locations, which seem to confer a further boost in performance.

For insertion predictions, we had two considerations: (i) inDelphi only predicts 1bp insertions; (ii) FORECasT groups all 1bp insertions that do not repeat the nucleotides flanking the cut site into one outcome. Thus, we compared only 1bp insertion frequencies and aggregated non-cut site-repeating insertions into one outcome. Both X-CRISP models performed comparably to Lindel, which was expected given that the X-CRISP insertion model is based on Lindel (Fig. 2). Moreover, X-CRISP/Lindel achieved lower JSDs than inDelphi, but higher than FORECasT (FORECasT|inDelphi mESC data X-CRISP KLD: 0.22|0.18, FORECasT: 0.21|0.17). We reason that since FORECasT is trained to predict the frequency of non-cut site-repeating insertions directly as a single group, it could have a slight advantage in these comparisons. In addition, the advantage of X-CRISP/Lindel over inDelphi could possibly be explained by the use of a wider sequence context surrounding the cut site (6bp vs. 3bp), providing additional degrees of freedom to model insertion frequencies.

### X-CRISP generalises well to frameshift prediction tasks

We further investigated if the detailed repair profiles predicted by the models could be useful to address higher-level prediction tasks, focusing on precision-*X*% targets and broader outcome categories. First, we assessed the prediction of precision-*X*% targets, defined as target sequences for which a single outcome accounts for at least *X*% of all reads assigned to any outcome (Shen *et al*., 2018). The ability to predict the precision-X% property can help with selecting CRISPR targets that maximise the proportion of the desired outcome relative to all other outcomes, for more precise gene editing. Performance was measured using precision (Table 4), recall, and Matthew’s correlation coefficient (Supplementary Table S4). On the FORECasT mESC test set, X-CRISP excelled in precision-20%, but came second to FORECAST in precision-30% through to precision-70% (Table 4). On the inDelphi mESC test set, FORECasT led in precision-20/30/40/50%, while X-CRISP MSE achieved top performance in precision-60%. Note that each model performed worse on inDelphi than on FORECasT data, highlighting the challenges of generalising to other datasets, even within the same cell type and genotype.

**Table 4.** Performance of six precision-X% prediction tasks, with *X* ∈ {20, 30, 40, 50, 60, 70}, measured using the precision performance score. All five models were trained on the 5900 FORECasT WT mESC target train set from Table 1, and then tested separately on the 3954 FORECasT and 1961 inDelphi WT mESC target test sets from Table 1.

| Test set | Model | Prediction of precision-$X\%$<br>(performance with $X = \dots$) | | | | | |
| --- | --- | --- | --- | --- | --- | --- | --- |
|  |  | 20 | 30 | 40 | 50 | 60 | 70 |
| FORECasT<br>WT mESC | X-CRISP KLD | **0.75** | 0.84 | 0.84 | 0.80 | 0.71 | 0.66 |
|  | X-CRISP MSE | **0.75** | 0.83 | 0.83 | 0.77 | 0.69 | 0.65 |
|  | FORECasT | 0.72 | **0.85** | **0.86** | **0.83** | **0.76** | **0.74** |
|  | Lindel | 0.71 | 0.77 | 0.69 | 0.60 | 0.33 | 0.33 |
|  | inDelphi | 0.40 | 0.45 | 0.24 | 0.09 | 0.06 | 0.03 |
| inDelphi<br>WT mESC | X-CRISP KLD | 0.73 | 0.70 | 0.57 | 0.40 | **0.28** | 0.00 |
|  | X-CRISP MSE | 0.73 | 0.70 | 0.54 | 0.34 | 0.21 | 0.00 |
|  | FORECasT | **0.77** | **0.83** | **0.71** | **0.46** | 0.21 | 0.00 |
|  | Lindel | 0.69 | 0.62 | 0.51 | 0.33 | 0.12 | 0.00 |
|  | inDelphi | 0.36 | 0.31 | 0.07 | 0.02 | 0.00 | 0.00 |

We further evaluated the performance of each model on six outcome profile aggregation tasks: deletion, 1bp insertion, 1bp deletion, 1bp frameshift, 2bp frameshift, and frameshift frequency prediction. We also included a CROTON model (Li *et al*., 2021) in the evaluation, which was originally developed to predict these six broader outcomes. To ensure comparability, we retrained CROTON on the same data as the other models, and aggregated the predictions of the other models per broader outcome.

The X-CRISP model achieved top or near-top performance across all tasks (Fig. 3 for MSE, Supplementary Fig. S23 for Pearson’s correlation). In both datasets, X-CRISP competed for the best deletion frequency and 1bp insertion frequency prediction performance with Lindel and CROTON, respectively. FORECasT led in 1bp deletion frequency prediction for both datasets, while X-CRISP excelled on all frameshift prediction tasks. Frameshift prediction is an especially important task, as frameshifts often result in gene knockouts, which are useful for studying gene function and developing therapeutics. We attribute the improved performance of X-CRISP on broader outcome prediction to the superiority already demonstrated when predicting detailed repair outcome profiles. In this case, the ability to predict more accurately across the entire frequency distribution seemed to translate into improved aggregate frequency predictions.

**Fig. 3:**
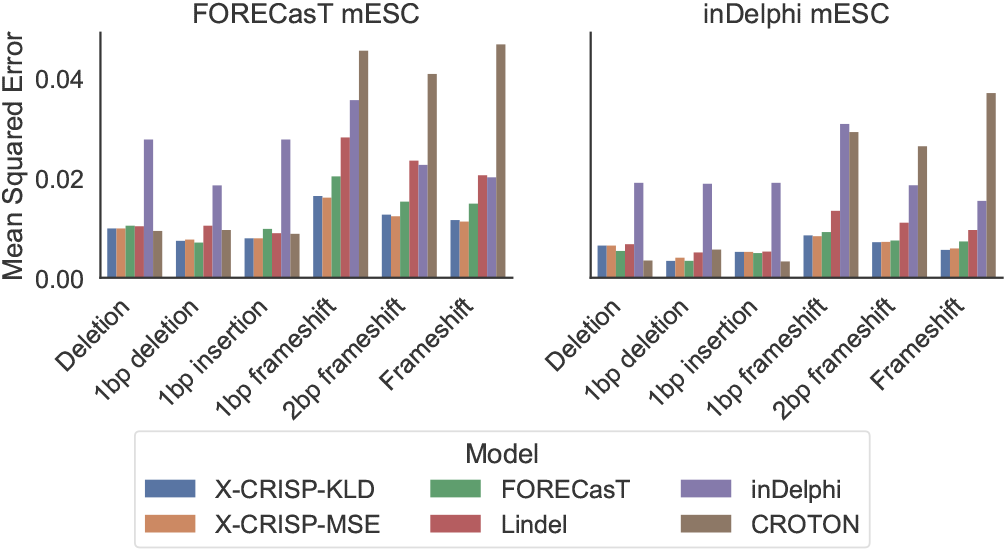
Broader repair outcome prediction performance. Mean-squared error (MSE) for six outcomes: deletion, 1bp insertion, 1bp deletion, 1bp frameshift, 2bp frameshift, and frameshift frequency prediction. Models trained on FORECasT WT mESC and tested on 3954 FORECasT and 1961 inDelphi WT mESC target sites.

Overall, the results indicate that repair outcome profiles predicted by X-CRISP generalise better than those predicted by other models to higher-level prediction tasks, such as precision-X% and broader outcome prediction. However, the loss function used to achieve optimal performance could be task-specific.

### Deletion prediction is most influenced by MH location

Compared to models like FORECasT and Lindel that break down (combinations of) outcome attributes such as MH length and position into thousands of categorical binary feature encodings for specific values or bins, X-CRISP preserves each of the five attributes it uses as a single feature in its original integer or real-valued range. For instance, Lindel splits MH length into five categorical bins (0, 1, 2, 3, and 4+) and couples them with positional and deletion length categories, resulting in a total of 2649 features where the effect of MH length alone cannot be easily discerned from the effects of other attributes. Similarly, FORECasT bins MH length into seven categories (“No MH”, 1, 2, 3, 4–6, 7–10, 11–15) and pairs those with additional attributes like deletion length or microhomology location, both also binned with varying ranges, yielding 525 MH-related features out of a total of 3633 used by the FORECasT model. In contrast, X-CRISP represents the MH length attribute using a single integer-valued feature and relies on the neural network to learn eventual interactions with other attributes or features in a data-driven manner. Similar observations can be made for the remaining attributes: where X-CRISP uses a single feature per attribute, FORECasT and Lindel typically use hundreds of features combining specific values or bins from multiple attributes.

While the encoding employed by FORECasT and Lindel is not necessarily limiting in terms of performance, given that the large numbers of features provide sufficient degrees of freedom to learn good prediction models, the dispersion and combination of attribute values across features make it challenging to isolate and interpret the contribution of each attribute on its own. On the other hand, the X-CRISP approach allows the learning of feature interactions and thus introduces black-box characteristics to the model. Nevertheless, post-hoc explainability tools such as SHAP are precisely designed to summarise and quantify how input attributes influence the predictions of (black-box) models, which we can readily use to gain insight into the impact of sequence characteristics on CRISPR outcome prediction across target sites and outcomes. To explore this, we obtained SHAP values for 400 randomly selected targets from the FORECasT test data to elucidate the influence of each feature on the predicted score of each X-CRISP submodel.

For MH-*based* deletions (Fig. 4A), the influence (SHAP value) of the left and right edges increased as they got closer to the cut site (feature values near zero), meaning that an MH-*based* deletion became more likely. We consider this intuitive since the edge features also denote MH location, and MHs closer to the cut site should be easier to select and anneal during repair due to their physical proximity. Counter-intuitively, as the gap between MHs decreased, the influence (SHAP value) of the gap feature decreased. We reason that this could be a correction applied by the model when both edges are near zero, to prevent the deletion frequency from growing disproportionately large. Longer MHs also led to larger MH-*based* deletion frequencies, yet were considerably less influential than MH location (left/right edge), while GC content exhibited minimal contribution. This could indicate that the selection of an MH during repair might be more influenced by MH position than by sequence content. For MH-*less* deletions, the gap and edge features showed similar behaviour to that described above for MH-*based* deletions, but with smaller SHAP value ranges.

**Fig. 4:**
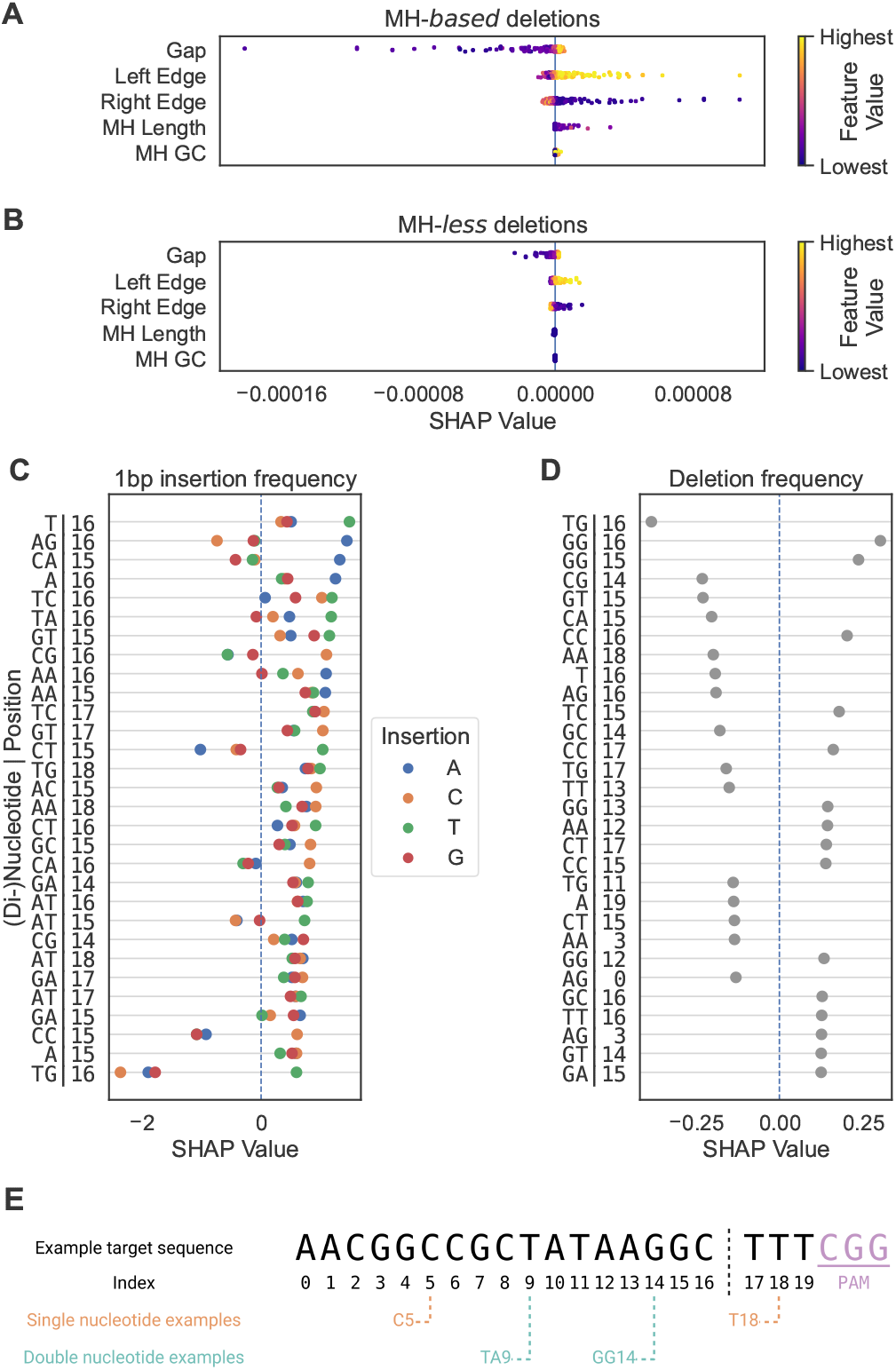
Influence of X-CRISP model features on repair outcome prediction. Feature effects expressed by SHAP values, calculated for 400 targets from the FORECasT WT mESC test set. Strip plots show SHAP values per feature for 10k randomly selected (A) MH-*based* and (B) MH-*less* deletions. (C) Top 30 features for all 1bp insertions, ranked by maximum SHAP value. (D) Top 30 features for the deletion-insertion model and ranked by absolute SHAP value, with SHAP values denoting impact on deletion frequency. (E) Indexing of single and dinucleotide features for an example target DNA sequence.

For insertions, we focused on the most prominent group: 1bp insertions (Fig. 4C). Cut site-proximal positions showed more influence than others, where an A or T immediately upstream of the DSB (at the 17th position or index 16 of the 20nt gRNA sequence, Fig. 4E for an illustration of target sequence indexing) promoted the insertion of an A or T, respectively. Insertions of C were positively influenced by the presence of C immediately upstream of, or CG centred at, the cut site. We did not observe strong associations between single or dinucleotide sequence content and G insertions. These findings align with existing literature (van Overbeek *et al*., 2016; Shen *et al*., 2018; Allen *et al*., 2019; Chen *et al*., 2019; Shi *et al*., 2019; Gisler *et al*., 2019).

For the deletion-insertion model, DSB-proximal nucleotides were strong influencers as well (Fig. 4D), with C or G at index 16 promoting deletions. However, A or T at index 16 (e.g. T|16, GT|15, CA|15, or AG|16) promoted insertions. A standout observation was that dinucleotide repeats centred at the cut site (except AA|16) favoured deletions of DSB-adjacent nucleotides, as previously observed elsewhere (Allen *et al*., 2019).

### Transfer learning greatly reduces data required for new domains

We explored if transfer learning could adapt pre-trained X-CRISP models to new domains, like other cell types or genomic contexts, and reduce the requirements of domain-specific training data.

We first examined changes in the distributions of repair outcomes between WT mESC cells and each of the four different target domains: human U2OS, human HAP1, NHEJ-deficient mESC, and human TREX2 cells (Fig. 5). The HAP1 cells revealed high similarity with mESC, exhibiting similar MH-*based* and MH-*less* deletion frequency distributions, average frequencies per deletion length, and insertion frequency distributions. The U2OS cells showed a larger variation in overall deletion frequency, a higher proportion of single A insertions, and a lower proportion of ≥3bp insertions. Cell lines with modified repair function deviated the most from the others: mESC *NHEJ*^*-/-*^ cells favoured MH-*based* deletions and led to less frequent insertions, especially 1bp insertions; TREX2 cells preferred deletions of 10-16bp over deletions of 3-9bp, and MH-*less* over MH-*based* deletions.

**Fig. 5:**
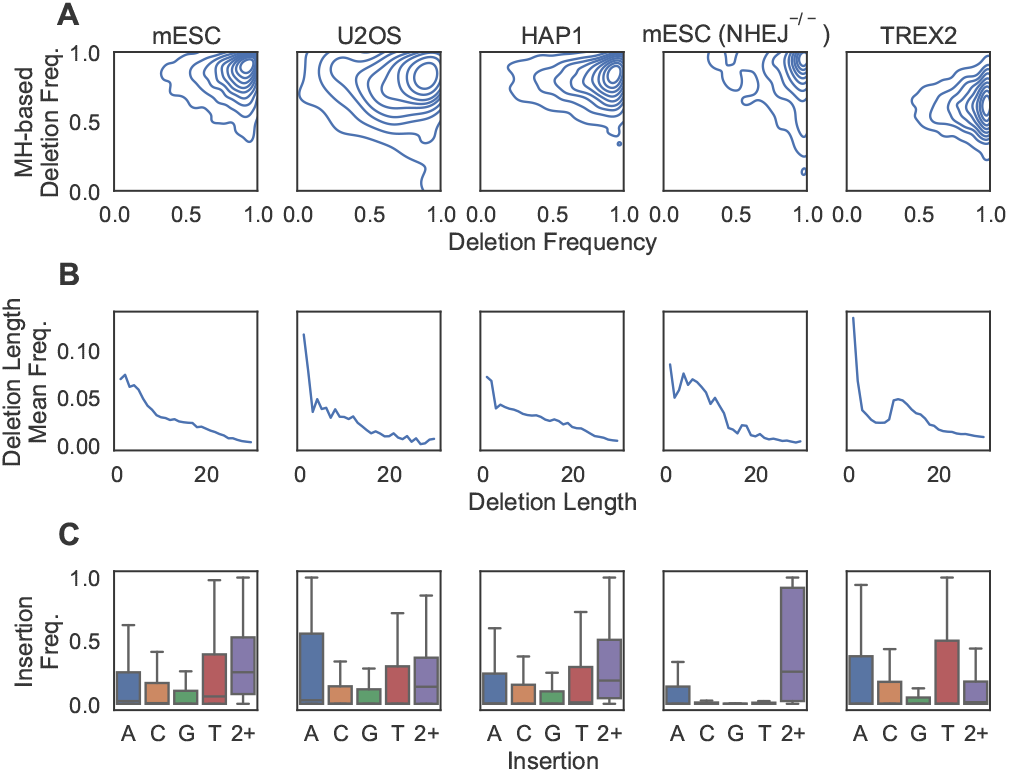
Repair outcome distributions across all X-CRISP processed repair outcome profiles observed for the mouse and human cell lines in Table 1: (A) densities for overall deletion frequency (horizontal) vs. MH-*based* deletion frequency (vertical), with each contour line denoting 10% of the data; (B) trend line of average frequency per deletion length; (C) frequency distributions of 1bp and ≥ 2bp insertions, outliers excluded.

For the transfer learning task, we adapted the X-CRISP models pre-trained on WT mESC data to each target domain using several techniques (Fig. 6 for JSD, Supplementary Fig. S24 for Pearson’s correlation). Each adapted model was further tested against held-out unseen data from the corresponding target domain. Pretraining on mESC data alone (SO, source only) generalised well to HAP1 cells (full repair profile mean JSD: mESC 0.426, HAP1 0.427), and achieved comparable performance to training directly on HAP1 data using at least 500 samples (TO, target only). The TL strategies showed little benefit here, likely due to the similarity between mESC and HAP1 (Fig. 5), requiring at least 500 target HAP1 samples before a consistent gain in performance could be observed across all TL methods (TL mean JSD: 0.420). For the transfer to U2OS cells, all TL methods significantly improved the full repair profile performance over the SO and TO baselines after retraining and fine-tuning on 50 target U2OS samples, with PF0 (retrained and fine-tuned on target data without layer freezing) achieving the best results (Wilcoxon two-tailed signed-rank test *p*-values PF0 vs. SO: 6 × 10^−92^; and vs. TO: 1 × 10^−137^). The improvement was also seen for both U2OS MH-*based* and MH-*less* deletion prediction. In addition, fine-tuning (FT) on U2OS cells improved deletion-insertion ratio prediction over SO and TO from only two target samples. These results indicate that TL could be used to reduce the number of target samples required to train new models for some target domains.

**Fig. 6:**
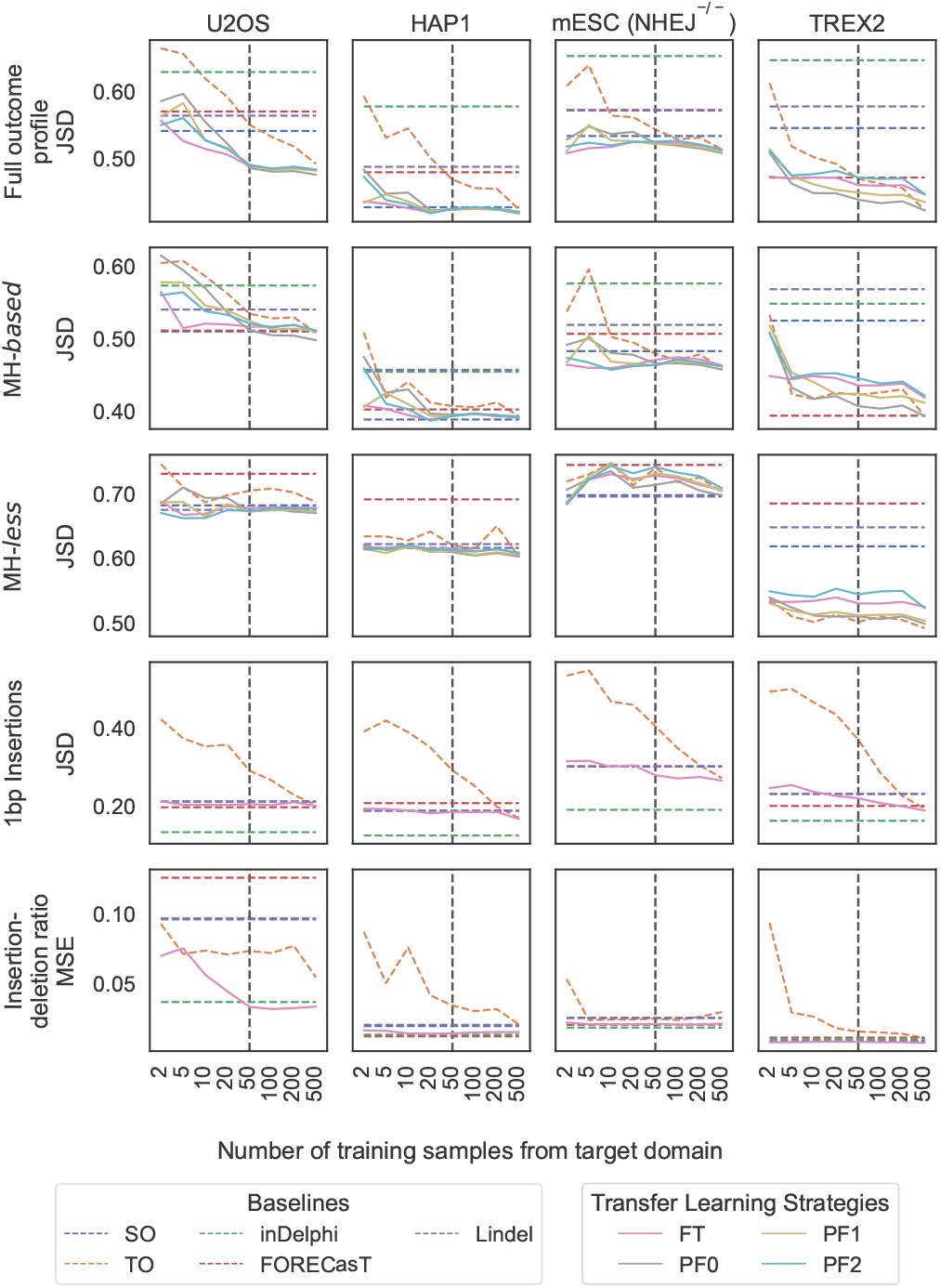
X-CRISP model adaptation to new domains or cell lines using transfer learning (TL). Prediction performance of baseline X-CRISP models and TL strategies, as the average JSD or the MSE between predicted and observed frequencies per model and number of training samples from the target domain. (Baseline models) TO, X-CRISP trained on target only; SO, X-CRISP trained on source only; FORECasT, Lindel, inDelphi. (Transfer learning) FT, X-CRISP pre-trained on source and fine-tuned for target; PF0-2, X-CRISP pre-trained on source and retrained + fine-tuned on target using 0-2 frozen hidden layers. (Top to bottom) Prediction models for full repair profile, MH-*based* deletions, MH-*less* deletions, 1bp insertions, and deletion-insertion ratios. Note: horizontal axis is categorical, thus it is not to scale; and the SO, FORECasT, inDelphi, and Lindel baselines do not use any samples from the target domain, so their performance remains constant along the horizontal axis. (Left to right) Models tested against U2OS (962), HAP1 (3950), TREX2 (3355), and mESC *NHEJ*^*-/-*^ (985) test target sites.

Lastly, we examined the results for cell lines with modified cellular DNA repair function. For mESC *NHEJ*^*-/-*^ cells, all TL methods showed small improvements in full repair profile performance over SO and TO after training on 50 target samples. Using 500 target samples, only PF0 outperformed SO and TO (mean JSD; PF0: 0.522, SO: 0.534), driven by small improvements in MH-*based* deletion prediction, consistent with the fact that impairment of NHEJ visibly altered MH-*based* deletion activity (Fig. 5). For TREX2 cells, PF0 was the most effective TL method, showing gains from just five target samples. Here, the insertion model displayed a significant performance benefit over both SO and TO baselines up to 500 target samples, while the deletion model did not benefit from TL, likely due to the large distribution shift towards longer deletions driven by the Cas9-fused three-prime exonuclease 2. The TO model achieved comparable performance to TL using 500 samples.

Overall, the most flexible transfer learning strategy (PF0, no layer freezing) showed the most effective adaptation to new domains, requiring only 50 target domain samples to consistently achieve results comparable or superior to training directly on a larger number of samples for the 4 different target domains. The gap between the least and most flexible TL strategies was especially apparent when adapting to larger changes between source and target context distributions (Fig.6, TREX2). These changes seemed largest when the underlying biological mechanisms for cutting or repairing the DNA were modified than across wild-type cell lines (Fig. 5, NHEJ^−*/*−^ and TREX2 vs. WT mESC, U2OS, and HAP1). This suggested a higher conservation of repair mechanisms between wild-type cells, even across mouse and human. Importantly, our results also showed that TL strategies provided the most benefit for genotypes affecting CRISPR-Cas9 function and DNA damage response (Fig. 6), creating opportunities to better understand the associated biological mechanisms and to improve the control over CRISPR-induced outcomes for more precise gene editing across fundamental and translational applications. Increasing the number of learnable parameters allowed the X-CRISP model to better realign to the larger changes in repair outcome distributions exhibited by genetically modified cells, highlighting that the effectiveness of TL is domain-dependent and considerations such as further tweaking of the models could be necessary for successful adaptation to more challenging contexts. We also analysed the impact of TL on the prediction performance of broader repair outcomes, where TL consistently improved the MSE over the baselines on the frameshift frequency tasks after 50 target domain samples (Supplementary Fig. S25, see Supplementary Fig. S26 for Pearson’s correlation). We envision that similar TL strategies could be used to adapt models for precise CRISPR therapeutic interventions considering the unique genetic landscape of specific patient cohorts or individual patients. To determine how successful such strategies could be would need extensive investigation and validation across a wide variety or at least a representative selection of human donors.

## Conclusions

We introduced X-CRISP, an interpretable and domain-adaptable model to predict the frequency of CRISPR repair outcomes. The X-CRISP model exhibited superior accuracy and generalisation in both detailed and broader repair outcome frequency prediction, especially when predicting frameshift mutations – a crucial task in experimental and therapeutic genome editing applications.

Top performance was achieved while retaining interpretability and flexibility to adapt X-CRISP models to additional genomic contexts. Contributing to this was the inclusion of informative features like deletion edges, which also function as indicators of MH location, and showed a prominent influence on deletion frequency prediction of X-CRISP models. Additionally, the ubiquitous representation of features for both MH-*based* and MH-*less* deletions, coupled with a non-linear neural network architecture, enabled X-CRISP to leverage a compact set of 5 features for improved and interpretable deletion prediction, compared to the next best models using ∼3k features.

Finally, we showed that pre-trained X-CRISP models could be successfully adapted using transfer learning for prediction in additional domains, spanning different organisms, cell lines, and DNA repair function characteristics. Transfer learning was effective from 50 target domain samples, suggesting that the typical range of thousands of domain-specific samples required to train a repair outcome prediction model for a new domain could be reduced by adapting existing pre-trained models, potentially using orders of magnitude less target domain samples. More flexible TL approaches, with freedom to adjust all weights of the prediction model, generally adapted better and were especially effective in transferring to more distant contexts characterised by larger changes in the repair outcome frequency distribution. These results highlight the potential of transfer learning to expedite the development of CRISPR repair outcome prediction models for contexts where generating extensive data may not be feasible and thus facilitate research involving CRISPR assays. Specifically, these models could help improve the design of guide RNAs to efficiently knock out specific genes with the aim of studying their function, or contribute to the development of CRISPR therapeutics by improving the design of guide RNAs to correct pathogenic variants across various genomic contexts.

## Acknowledgments

The authors acknowledge the High-Performance Compute cluster of the INSY department at Delft University of Technology, and early access to SIQ kindly provided by Marcel Tijsterman and Robin van Schendel from the Leiden University Medical Centre.

## References

Adli, M. (2018). The CRISPR tool kit for genome editing and beyond. Nature Communications, 9(1), 1–13.

Allen, F., et al. (2019). Predicting the mutations generated by repair of Cas9-induced double-strand breaks. Nature Biotechnology, 37(1), 64–72.

Bhattacharya, D., et al. (2015). CRISPR/Cas9: An inexpensive, efficient loss of function tool to screen human disease genes in Xenopus. Developmental Biology, 408(2), 196–204.

Cahill, D., et al. (2006). Mechanisms of eukaryotic DNA double-strand break repair. Frontiers in Bioscience (Landmark Ed), 11(2), 1958–1976.

Chehelgerdi, M., et al. (2024). Comprehensive review of crispr-based gene editing: mechanisms, challenges, and applications in cancer therapy. Molecular cancer, 23(1), 9.

Chen, W., et al. (2019). Massively parallel profiling and predictive modeling of the outcomes of CRISPR/Cas9-mediated double-strand break repair. Nucleic Acids Research, 47(15), 7989–8003.

[dataset]* Allen, F., et al. (2018). Predicting the mutations generated by repair of Cas9-induced double-strand breaks. European Nucleotide Archive. PRJEB29746.

[dataset]* Shen, M. W., et al. (2018). Deep sequencing of Cas9 editing outcomes in mouse cells. NCBI Sequence Read Archive. SRP141144.

Gisler, S., et al. (2019). Multiplexed Cas9 targeting reveals genomic location effects and grna-based staggered breaks influencing mutation efficiency. Nature Communications, 10(1), 1598.

Hsu, P. D., et al. (2014). Development and applications of CRISPR-Cas9 for genome engineering. Cell, 157(6), 1262–1278.

Jinek, M., et al. (2012). A programmable dual-RNA-guided DNA endonuclease in adaptive bacterial immunity. Science, 337(6096), 816–821.

Kingma, D. P. and Ba, J. (2014). Adam: A method for stochastic optimization. arXiv preprint 1412.6980.

Koike-Yusa, H., et al. (2014). Genome-wide recessive genetic screening in mammalian cells with a lentiviral CRISPR-guide RNA library. Nature Biotechnology, 32(3), 267–273.

Kullback, S. and Leibler, R. A. (1951). On information and sufficiency. The Annals of Mathematical Statistics, 22(1), 79–86.

Leenay, R. T., et al. (2019). Large dataset enables prediction of repair after CRISPR–Cas9 editing in primary T cells. Nature Biotechnology, 37(9), 1034–1037.

Lehmann, A. R. and Taylor, E. M. (2001). Conservation of eukaryotic DNA repair mechanisms. In DNA Damage and Repair, pages 377–401. Springer.

Leinonen, R., et al. (2010). The European Nucleotide Archive. Nucleic Acids Research, 39(suppl 1), D28–D31.

Li, V. R., et al. (2021). CROTON: an automated and variant-aware deep learning framework for predicting CRISPR/Cas9 editing outcomes. Bioinformatics, 37(Supplement 1), i342–i348.

Lundberg, S. M. and Lee, S.-I. (2017). A unified approach to interpreting model predictions. pages 4765–4774.

Maxmen, A. (2019). Faster, better, cheaper: the rise of CRISPR in disease detection. Nature, 566(7745), 437–438.

Naert, T., et al. (2020). Maximizing CRISPR/Cas9 phenotype penetrance applying predictive modeling of editing outcomes in Xenopus and zebrafish embryos. Scientific Reports, 10(1), 1–12.

Needleman, S. B. and Wunsch, C. D. (1970). A general method applicable to the search for similarities in the amino acid sequence of two proteins. Journal of molecular biology, 48(3), 443–453.

Pan, S. J. and Yang, Q. (2009). A survey on transfer learning. IEEE Transactions on Knowledge and Data Engineering, 22(10), 1345–1359.

Scully, R., et al. (2019). DNA double-strand break repair-pathway choice in somatic mammalian cells. Nature Reviews Molecular Cell Biology, 20(11), 698–714.

Shen, M. W., et al. (2018). Predictable and precise template-free CRISPR editing of pathogenic variants. Nature, 563(7733), 646–651.

Shi, X., et al. (2019). Cas9 has no exonuclease activity resulting in staggered cleavage with overhangs and predictable di-and tri-nucleotide CRISPR insertions without template donor. Cell Discovery, 5(1), 1–4.

Smith, T. F., et al. (1981). Identification of common molecular subsequences. Journal of molecular biology, 147(1), 195–197.

Tan, C., et al. (2018). A survey on deep transfer learning. In International Conference on Artificial Neural Networks, pages 270–279. Springer.

van Overbeek, M., et al. (2016). DNA repair profiling reveals nonrandom outcomes at Cas9-mediated breaks. Molecular Cell, 63(4), 633–646.

van Schendel, R., et al. (2022). SIQ: easy quantitative measurement of mutation profiles in sequencing data. NAR Genomics and Bioinformatics, 4(3), lqac063.

Wang, T., et al. (2014). Genetic screens in human cells using the CRISPR-Cas9 system. Science, 343(6166), 80–84.

Weiss, K., et al. (2016). A survey of transfer learning. Journal of Big Data, 3(1), 1–40.

Yue, X., et al. (2020). DNA-PKcs: A multi-faceted player in DNA damage response. Frontiers in Genetics, page 1692.

Zhang, J., et al. (2014). PEAR: a fast and accurate Illumina Paired-End reAd mergeR. Bioinformatics, 30(5), 614–620.

