## Supplementary Materials for "X-CRISP: Domain-Adaptable and Interpretable CRISPR Repair Outcome Prediction"

S26 X-CRISP model adaptation via Transfer Learning on aggregate tasks evaluated via Pearson’s correlation 18

### Supplementary Tables

| Source | Cell Type | Genotype | Sample Accession(s) |
|---|---|---|---|
| FORECasT [1] | mESC | Wild-Type | SAMEA5093999, SAMEA5094000 |
| | K562 | TREX2 | SAMEA104549464 |
| | HAP1 | Wild-Type | SAMEA5094017, SAMEA5094018 |
| inDelphi [4] | mESC | Wild-Type | SAMN08955971 |
| | U2OS | Wild-Type | SAMN09689449 |
| | mESC | Prkdc$^{-/-}$Lig4$^{-/-}$ | SAMN08955971 |

Table S1: CRISPR repair outcome sequencing data accession numbers.

| Model | $\beta_1$ | $\beta_2$ | Learning rate (LR) | LR decay ($\gamma$) | Penalty (optimised via 5-fold CV) | Penalty (final) |
|---|---|---|---|---|---|---|
| Training on source domain | | | | | | |
| Deletion | .99 | .999 | 0.05 | 0.999 | Tested L2 regularisation with weights in $\{0.01, 0.001, 0.0001, 0.00001, 0.000001\}$ | 0.00001 (L2) |
| Insertion | .99 | .999 | 0.001 | 0.99 | Tested L1 and L2 regularisation independently with weights in the range of between $10^{-10}$ and $10^{-1}$ | 0.0005011872 (L1) |
| Deletion-insertion | .99 | .999 | 0.001 | 0.99 | Tested L1 and L2 regularisation independently with weights in the range of between $10^{-10}$ and $10^{-1}$ | 0.00025118864 (L1) |
| Transfer Learning: Further training on target domain | | | | | | |
| Deletion | .99 | .999 | 0.05 | 0.999 | NA | 0.000001 (L2) |
| Insertion | NA | NA | NA | NA | NA | NA |
| Deletion-insertion | NA | NA | NA | NA | NA | NA |
| Transfer Learning: Fine-tuning on target domain | | | | | | |
| Deletion | .99 | .999 | 0.0005 | 0.999 | NA | 0.000001 (L2) |
| Insertion | .99 | .999 | 0.001 | 0.99 | NA | 0.0005011872 (L1) |
| Deletion-insertion | .99 | .999 | 0.0001 | 0.99 | NA | 0.00025118864 (L1) |

Table S2: Hyperparameters for model training on source domain and transfer to target domains.

| Model | Input | Model architecture | Predicts frequency of... |
|---|---|---|---|
| FORECasT [1] | 3,633 binary features describing each potential indel | Logistic regression model | All deletion outcomes touching the nucleotide directly downstream of the cut site, up a deletion length of 30nt, and all insertions of up to 2nt in the -3/0 window upstream of the cut site. |
| Lindel [2] | 2,649 binary features describing MH locations within the target sequence and 384 binary features for the one-hot encoded target sequence | Three multioutput logistic regression models | All 536 of 550 possible deletion outcomes of length <30nt that overlap with the -3/+2 window around the cut site, 21 insertion outcomes including all single and dinucleotide insertions and insertions of length $\geq$ 3bp. |
| inDelphi [4] | 3 features describing each MH deletion, one-hot encodings of the -3, -4, and -5 nucleotides (upstream of the PAM), and 2 features describing the predicted distribution of deletions | Combined two neural networks and k-nearest neighbour model | All MH-based deletions and all non-MH-based deletions grouped by deletion length, with a deletion length < 60nt, and all single nucleotide insertions. |
| | | | Predicts ratio of... |
| CROTON [3] | One-hot encoded 60nt target sequence | Convolutional neural network | Aggregate categories of outcomes against all others. Six separate CROTON models trained for predictions of: deletions, 1nt insertions, 1nt deletions, 1nt frameshift mutations, 2nt frameshift mutations, frameshift mutations. |

Table S3: Repair outcome prediction model comparison. FORECasT, Lindel, and inDelphi predict repair outcome profiles. CROTON predicts the ratio of six aggregate categories of repair outcomes against all others.

| Test set | Model | | Performance of six precision-X% prediction tasks (prediction: is the target precision-$X$%? yes/no) | | | | | |
|---|---|---|---|---|---|---|---|---|
| | | $X =$ | 20 | 30 | 40 | 50 | 60 | 70 |
| | | | Precision performance score | | | | | |
| FORECasT WT mESC [1] | X-CRISP KLD | | **0.746** | 0.841 | 0.844 | 0.795 | 0.702 | 0.655 |
| | X-CRISP MSE | | **0.751** | 0.831 | 0.828 | 0.766 | 0.695 | 0.653 |
| | FORECasT | | 0.723 | **0.853** | **0.858** | **0.836** | **0.755** | **0.739** |
| | Lindel | | 0.714 | 0.769 | 0.693 | 0.596 | 0.333 | 0.333 |
| | inDelphi | | 0.400 | 0.447 | 0.238 | 0.086 | 0.059 | 0.029 |
| inDelphi WT mESC [4] | X-CRISP KLD | | 0.726 | 0.702 | 0.569 | 0.396 | **0.278** | 0.000 |
| | X-CRISP MSE | | 0.728 | 0.698 | 0.537 | 0.340 | 0.211 | 0.000 |
| | FORECasT | | **0.771** | **0.827** | **0.704** | **0.464** | 0.211 | 0.000 |
| | Lindel | | 0.682 | 0.617 | 0.512 | 0.3326 | 0.115 | 0.000 |
| | inDelphi | | 0.363 | 0.313 | 0.071 | 0.022 | 0.000 | 0.000 |
| | | | Recall performance score | | | | | |
| FORECasT WT mESC [1] | X-CRISP KLD | | **0.643** | 0.597 | 0.474 | 0.412 | 0.301 | 0.191 |
| | X-CRISP MSE | | 0.652 | **0.617** | **0.518** | **0.458** | **0.378** | **0.250** |
| | FORECasT | | 0.522 | 0.428 | 0.323 | 0.244 | 0.157 | 0.080 |
| | Lindel | | 0.520 | 0.329 | 0.172 | 0.075 | 0.011 | 0.004 |
| | inDelphi | | 0.559 | 0.368 | 0.280 | 0.173 | 0.154 | 0.093 |
| inDelphi WT mESC [4] | X-CRISP KLD | | 0.712 | 0.771 | 0.669 | 0.462 | 0.333 | 0.000 |
| | X-CRISP MSE | | **0.726** | **0.780** | **0.706** | **0.555** | **0.400** | 0.000 |
| | FORECasT | | 0.613 | 0.572 | 0.460 | 0.359 | 0.216 | 0.000 |
| | Lindel | | 0.648 | 0.551 | 0.412 | 0.271 | 0.162 | 0.000 |
| | inDelphi | | 0.608 | 0.316 | 0.075 | 0.020 | 0.000 | 0.000 |
| | | | Matthew's correlation coefficient (MCC) | | | | | |
| FORECasT WT mESC [1] | X-CRISP KLD | | 0.283 | 0.498 | 0.515 | 0.504 | 0.418 | 0.337 |
| | X-CRISP MSE | | **0.298** | **0.501** | **0.534** | **0.519** | **0.468** | **0.386** |
| | FORECasT | | 0.172 | 0.384 | 0.409 | 0.388 | 0.311 | 0.232 |
| | Lindel | | 0.150 | 0.267 | 0.218 | 0.143 | 0.039 | 0.042 |
| | inDelphi | | -0.016 | 0.143 | 0.095 | 0.020 | 0.036 | 0.027 |
| inDelphi WT mESC [4] | X-CRISP KLD | | 0.370 | 0.589 | **0.535** | 0.387 | **0.292** | **-0.001** |
| | X-CRISP MSE | | **0.381** | **0.591** | 0.528 | **0.388** | 0.275 | -0.002 |
| | FORECasT | | 0.376 | 0.561 | 0.496 | 0.367 | 0.198 | -0.003 |
| | Lindel | | 0.253 | 0.387 | 0.365 | 0.268 | 0.146 | -0.004 |
| | inDelphi | | 0.055 | 0.166 | 0.010 | -0.003 | -0.006 | 0.000 |

Table S4: Performance of five models for six different precision-X% prediction tasks and two test sets, according to three performance metrics. The goal of each prediction task was to predict if a given target had a precision-X% outcome or not for a specific value of X. We evaluated six precision-X% prediction tasks, each for a different value of $X \in \{20, 30, 40, 50, 60, 70\}$, using three performance metrics: precision, recall, and Matthew's correlation coefficient. All models were trained on the FORECasT wild-type mESC train set and were then tested separately on the FORECast and the inDelphi wild-type mESC test sets from Table 1 of the main article, as specified in the "Test set" column. Note that precision-X% refers to a target sequence for which a single CRISPR-induced repair outcome accounts for at least $X$% of all reads assigned to any outcome observed for that target sequence, as defined by Shen et al. in [4].

#### Supplementary Figures

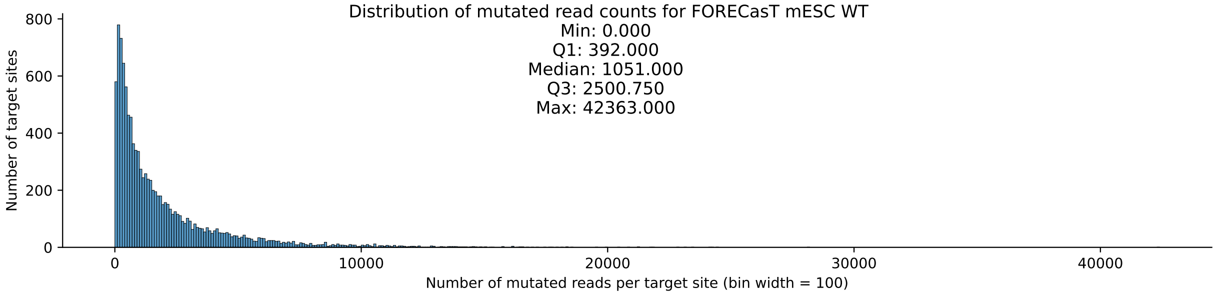

Figure S1: Distribution of mutated read counts per target site for the FORECasT mESC WT data. The horizontal axis shows the number of mutated reads assigned to each target site in bins of 100 in width. The vertical axis shows the number of target sites within each bin.

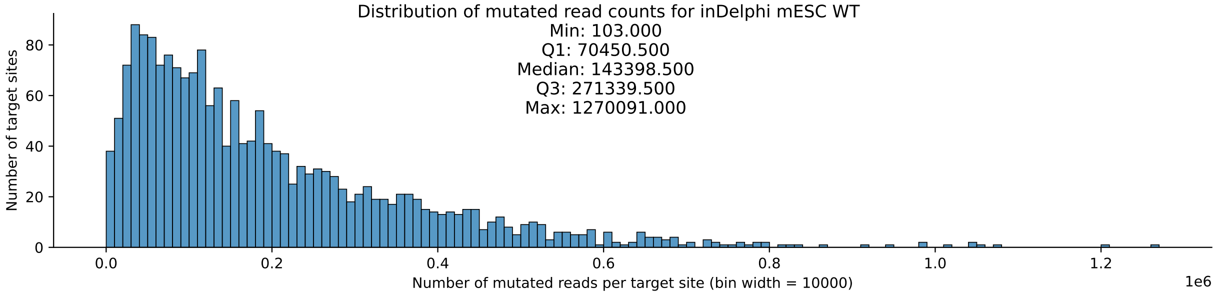

Figure S2: Distribution of mutated read counts per target site for the inDelphi mESC WT data. The horizontal axis shows the number of mutated reads assigned to each target site in bins of 10000 in width. The vertical axis shows the number of target sites within each bin.

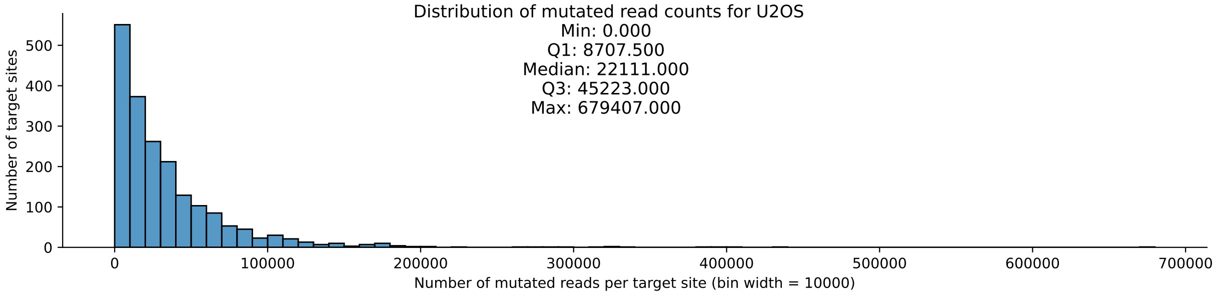

Figure S3: Distribution of mutated read counts per target site for the U2OS data. The horizontal axis shows the number of mutated reads assigned to each target site in bins of 10000 in width. The vertical axis shows the number of target sites within each bin.

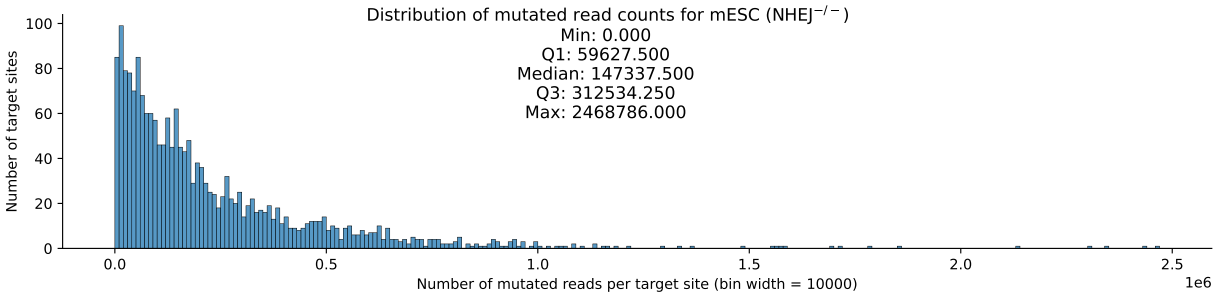

Figure S4: Distribution of mutated read counts per target site for the NHEJ-deficient mESC data. The horizontal axis shows the number of mutated reads assigned to each target site in bins of 10000 in width. The vertical axis shows the number of target sites within each bin.

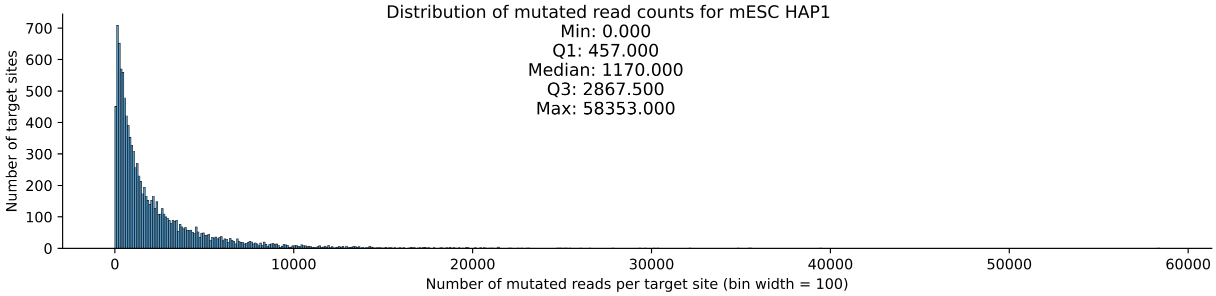

Figure S5: Distribution of mutated read counts per target site for the HAP1 data. The horizontal axis shows the number of mutated reads assigned to each target site in bins of 100 in width. The vertical axis shows the number of target sites within each bin.

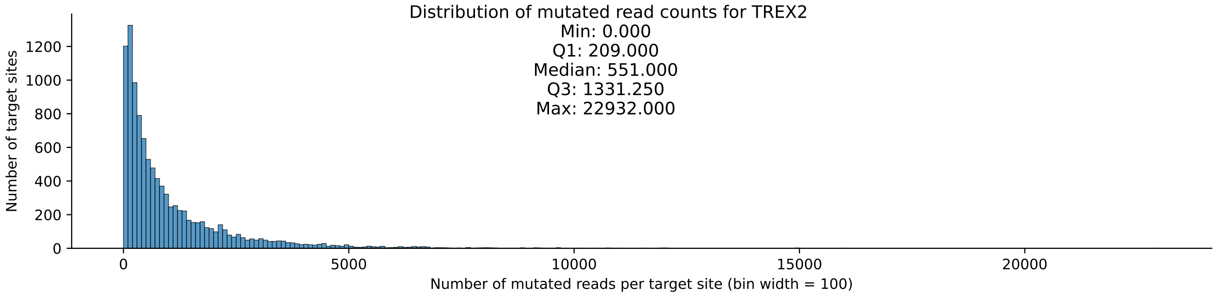

Figure S6: Distribution of mutated read counts per target site for the TREX2 data. The horizontal axis shows the number of mutated reads assigned to each target site in bins of 100 in width. The vertical axis shows the number of target sites within each bin.

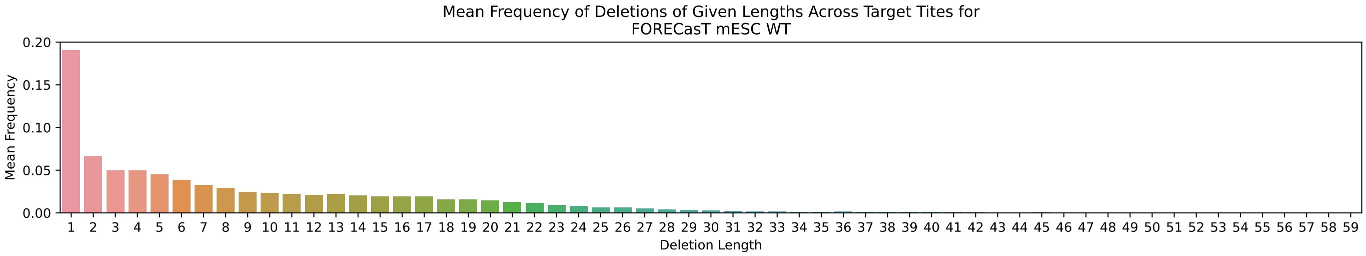

Figure S7: Mean frequency of deletions of given lengths across target sites for the FORECasT mESC WT data.

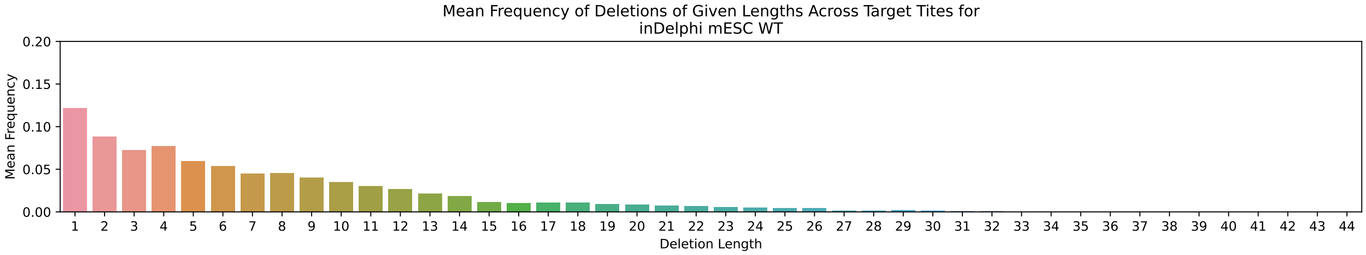

Figure S8: Mean frequency of deletions of given lengths across target sites for the inDelphi mESC WT data.

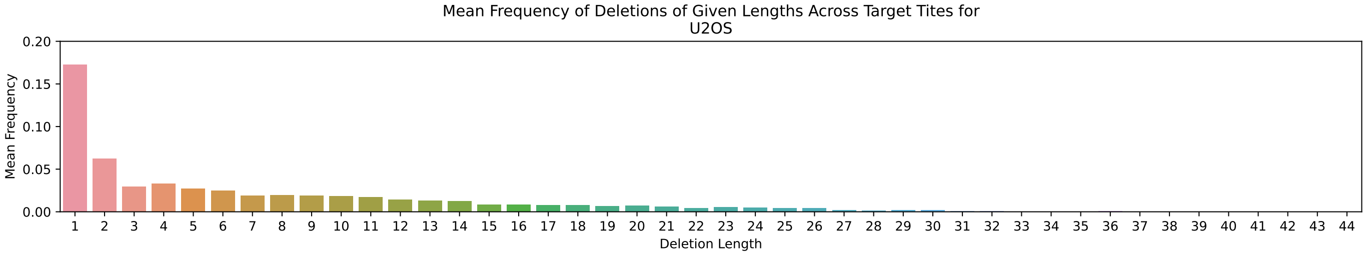

Figure S9: Mean frequency of deletions of given lengths across target sites for the U2OS data.

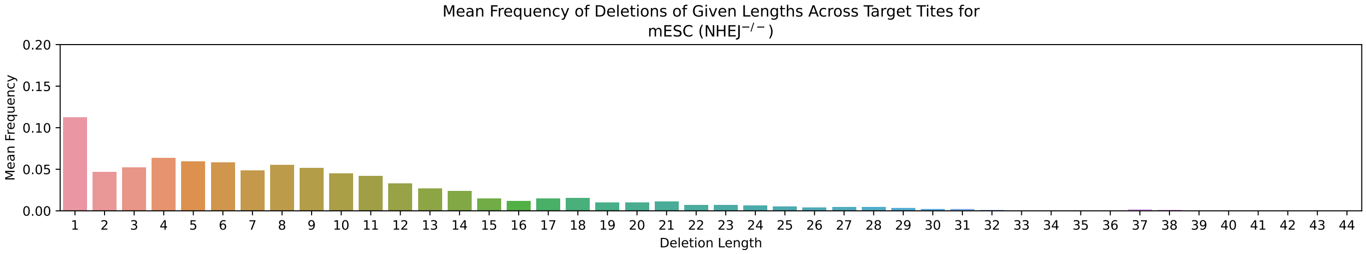

Figure S10: Mean frequency of deletions of given lengths across target sites for the NHEJ-deficient mESC data.

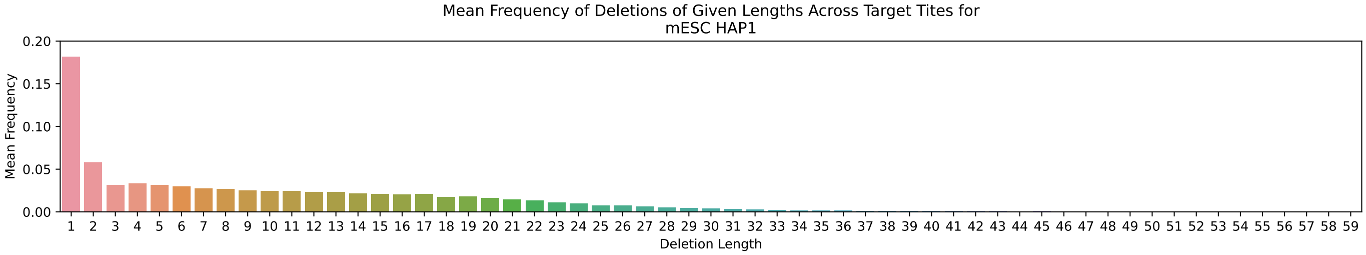

Figure S11: Mean frequency of deletions of given lengths across target sites for the HAP1 data.

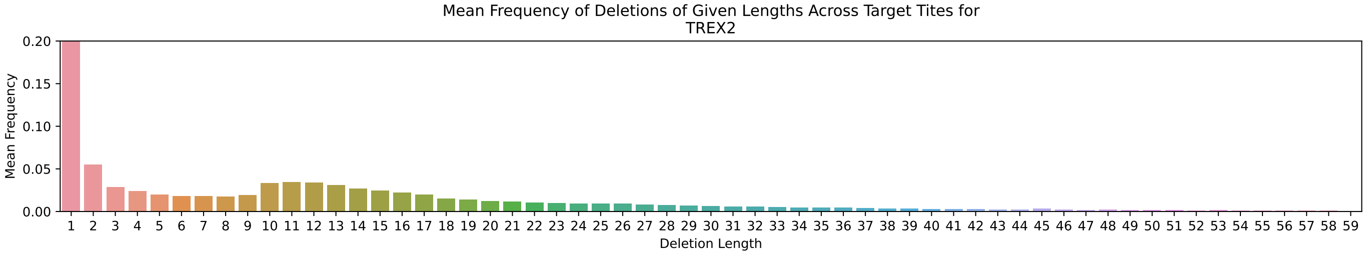

Figure S12: Mean frequency of deletions of given lengths across target sites for the TREX2 data.

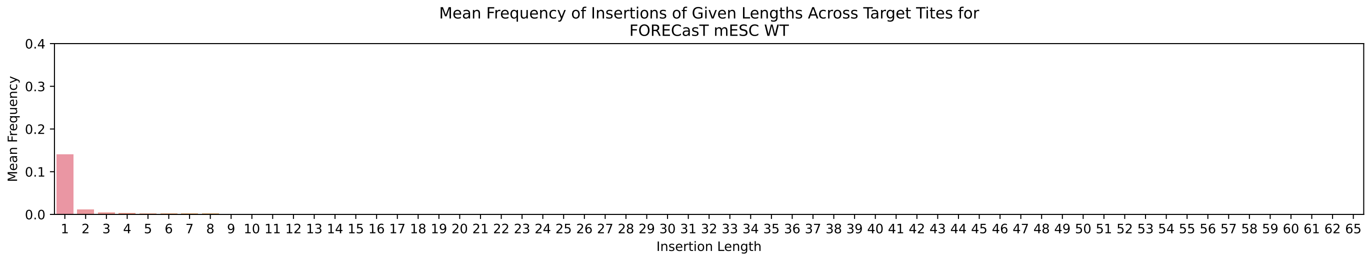

Figure S13: Mean frequency of insertions of given lengths across target sites for the FORECasT mESC WT data.

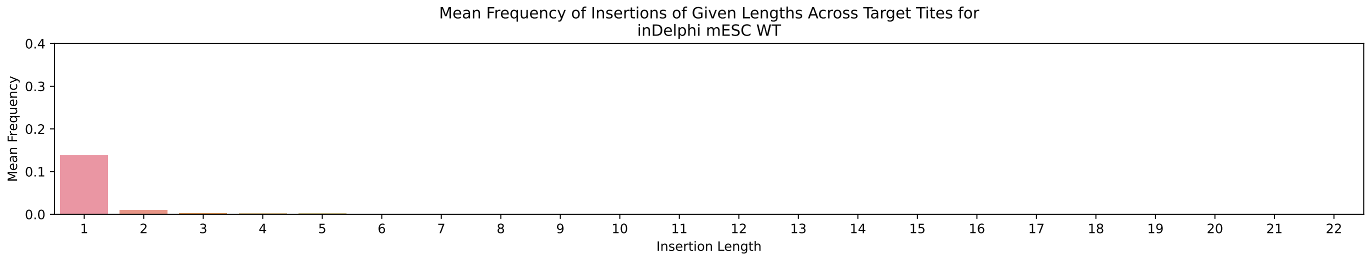

Figure S14: Mean frequency of insertions of given lengths across target sites for the inDelphi mESC WT data.

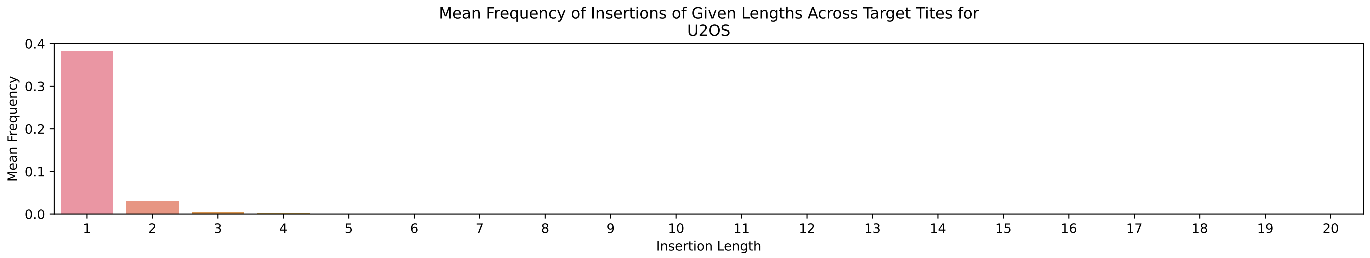

Figure S15: Mean frequency of insertions of given lengths across target sites for the U2OS data.

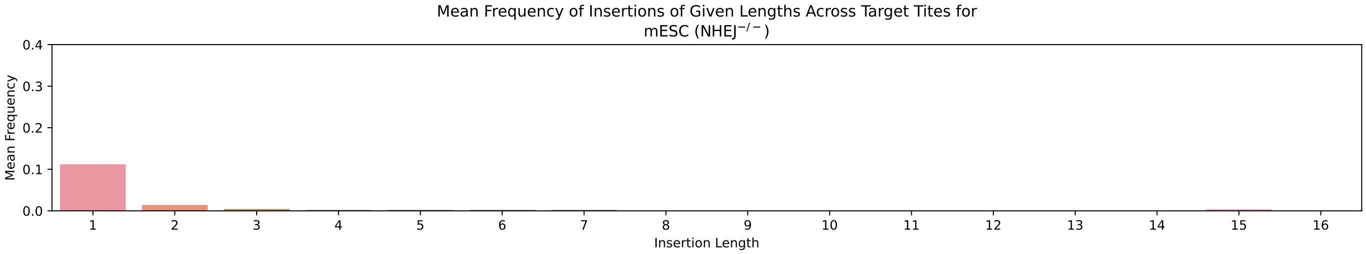

Figure S16: Mean frequency of insertions of given lengths across target sites for the NHEJ-deficient mESC data.

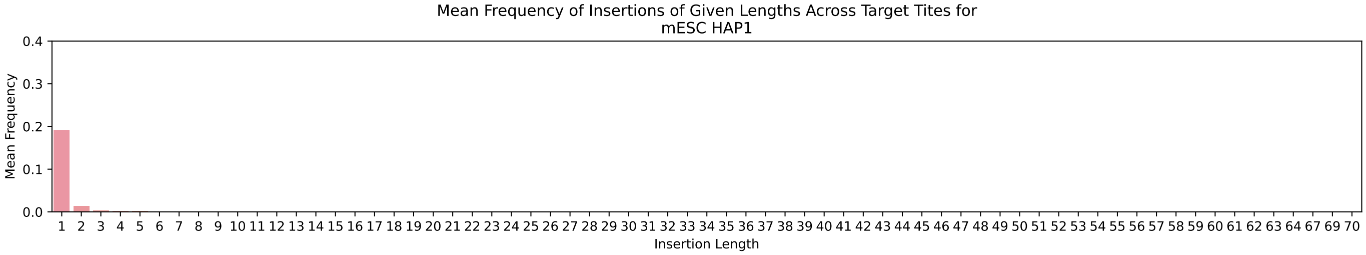

Figure S17: Mean frequency of insertions of given lengths across target sites for the HAP1 data.

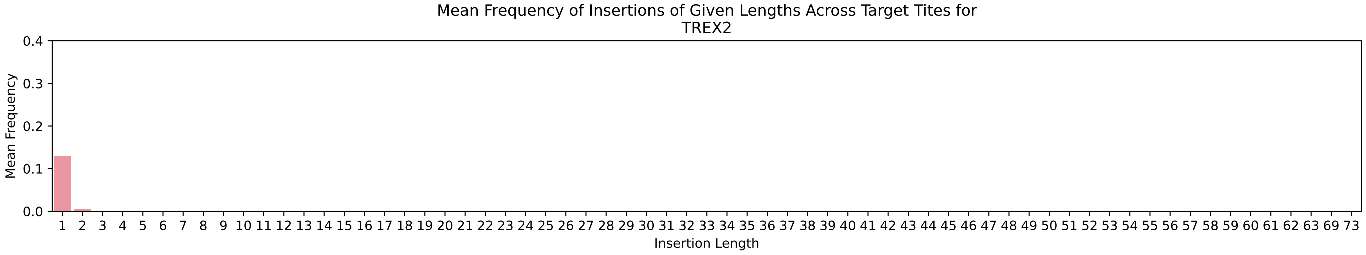

Figure S18: Mean frequency of insertions of given lengths across target sites for the TREX2 data.

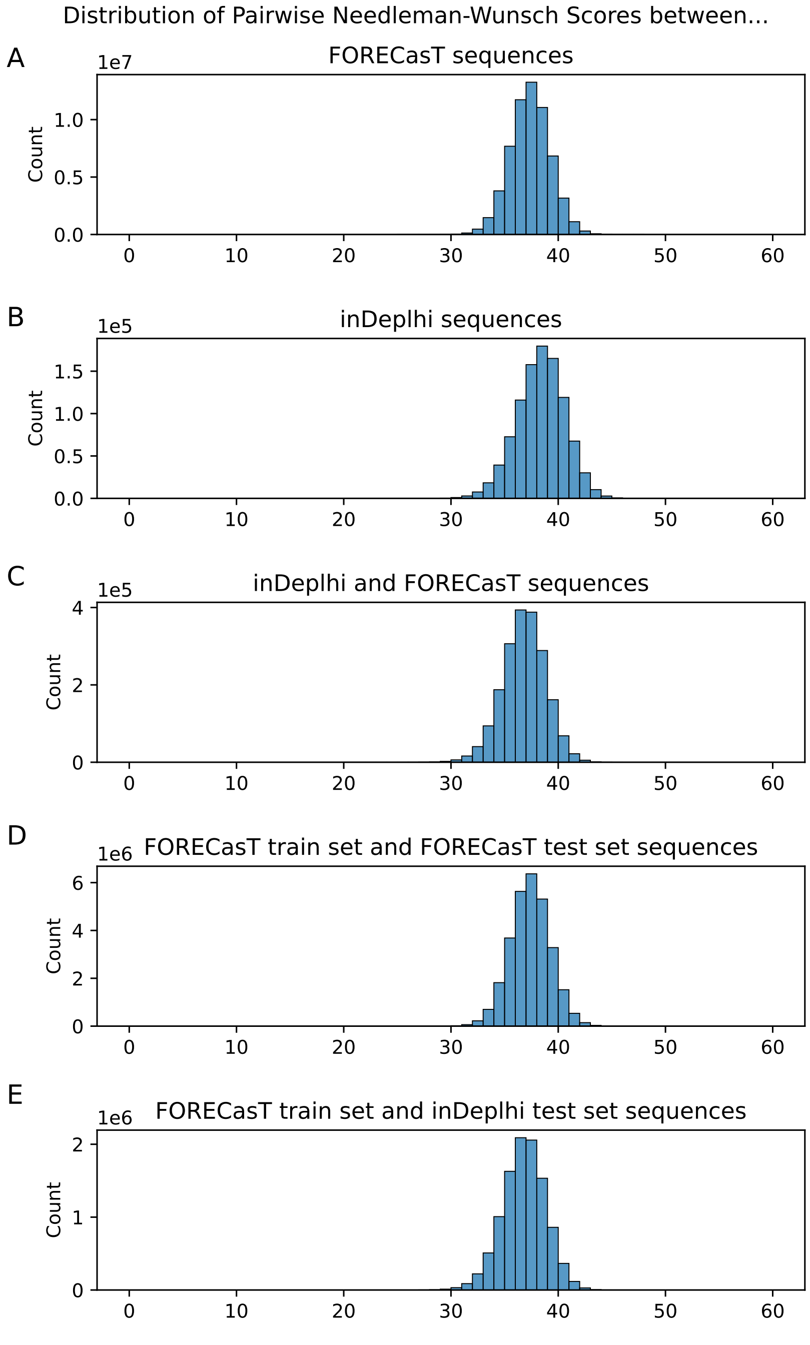

Figure S19: Pairwise Needleman-Wunsch scores between (A) sequences in the FORECasT dataset (B) sequences from the inDelphi dataset (C) sequences from the inDelphi dataset and the FORECasT dataset (D) sequences from the inDelphi dataset and the FORECasT dataset (E) sequences from the FORECasT train set and the inDelphi test set. All sequences are 60bp in length, centred at the cut site. Scoring parameters used: match: 1.0, mismatch: 0.0, gap: 0.0.

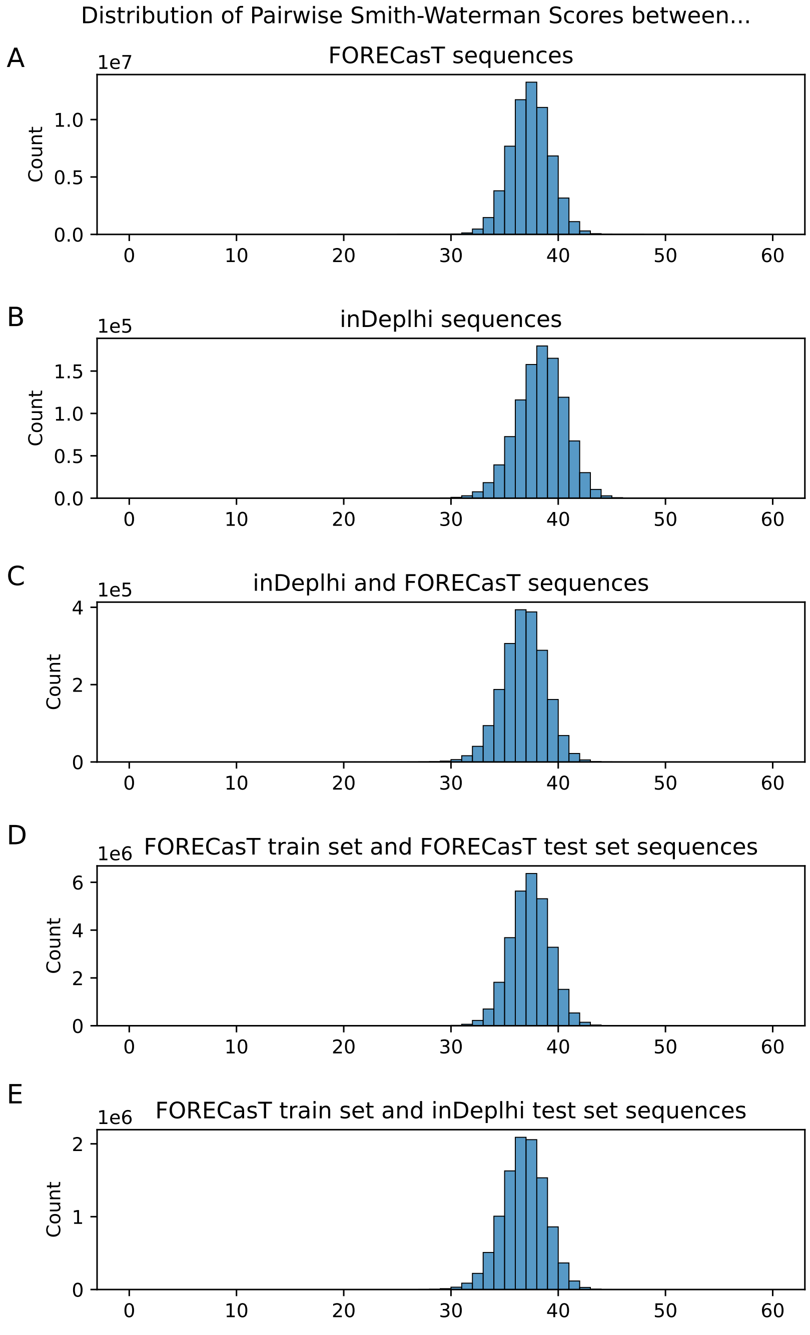

Figure S20: Pairwise Smith-Waterman scores between (A) sequences in the FORECasT dataset (B) sequences from the inDelphi dataset (C) sequences from the inDelphi dataset and the FORECasT dataset (D) sequences from the inDelphi dataset and the FORECasT dataset (E) sequences from the FORECasT train set and the inDelphi test set. All sequences are 60bp in length, centred at the cut site. Scoring parameters used: match: 1.0, mismatch: 0.0, gap: 0.0.

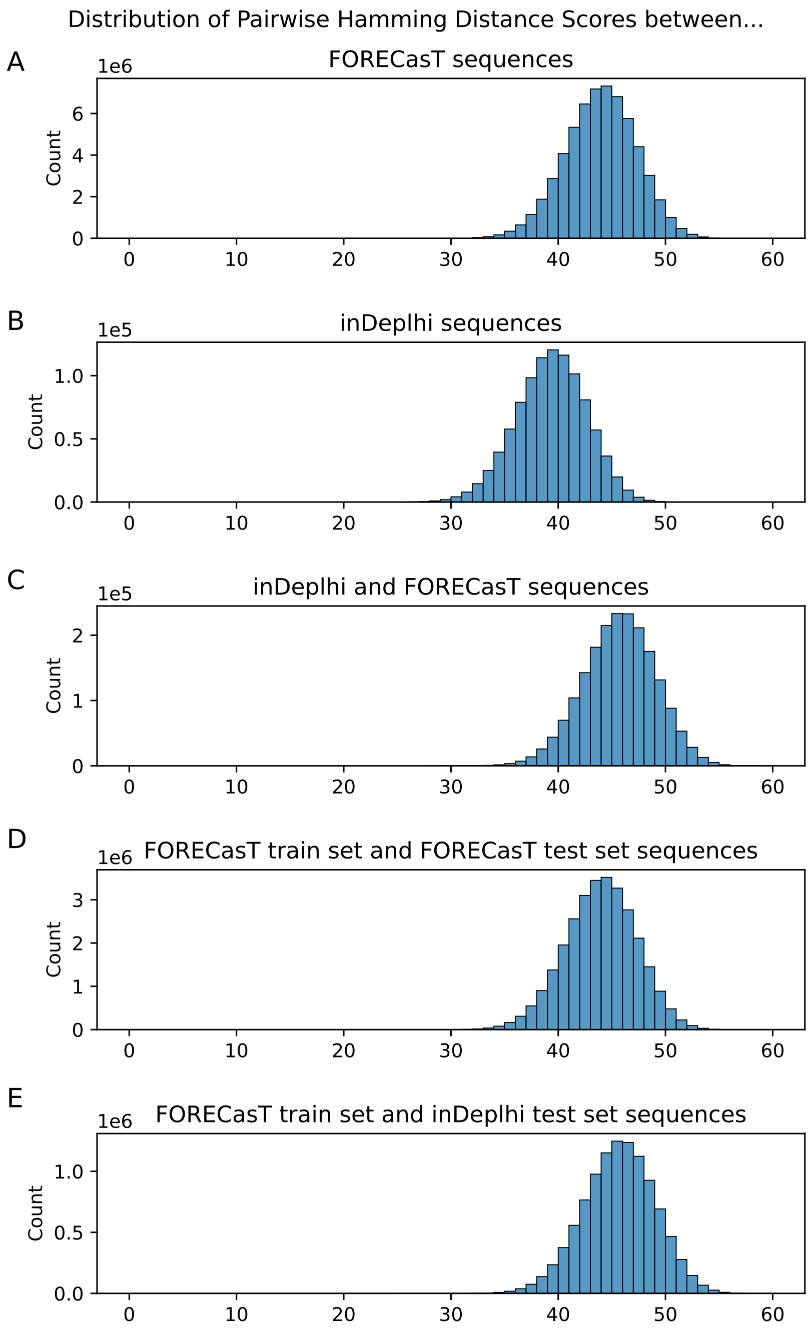

Figure S21: Pairwise Hamming distances between (A) sequences in the FORECasT dataset (B) sequences from the inDelphi dataset (C) sequences from the inDelphi dataset and the FORECasT dataset (D) sequences from the inDelphi dataset and the FORECasT dataset (E) sequences from the FORECasT train set and the inDelphi test set. All sequences are 60bp in length, centred at the cut site.

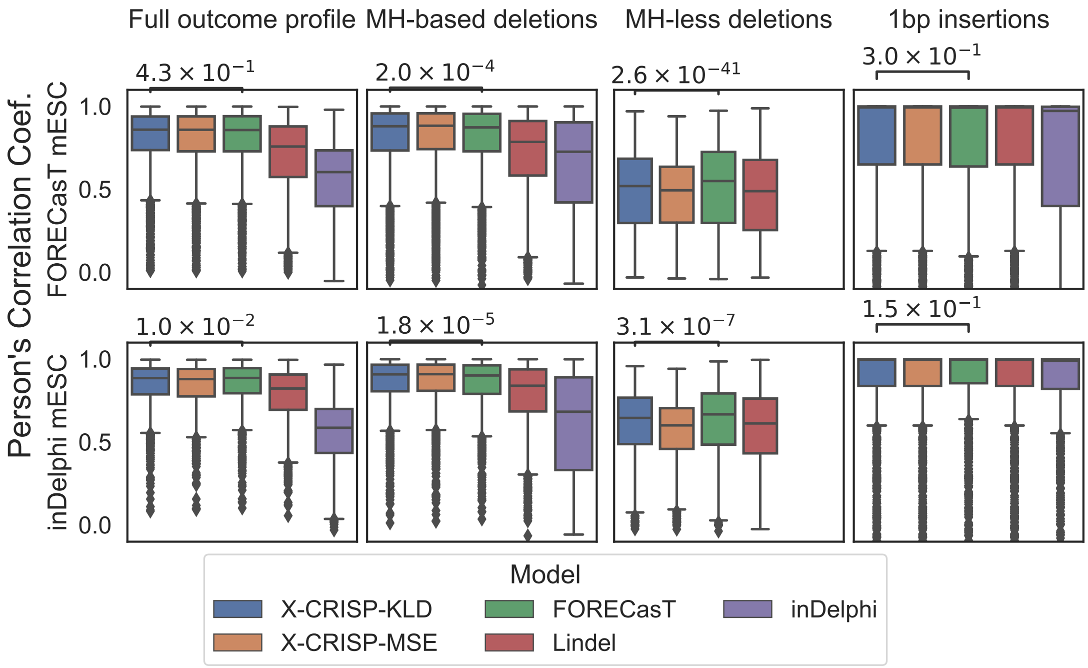

Figure S22: Detailed repair outcome prediction performance. Pearson's correlation coefficient between predicted and observed outcomes for (top) FORECasT or (bottom) inDelphi test data, considering: (left to right) original publication outcomes; common MH-*based* deletions; common MH-*less* deletions; 1bp insertions. Significance $p$-values calculated using Wilcoxon signed-rank tests, comparing X-CRISP KLD to the best of the non-X-CRISP models. For 1bp insertions, X-CRISP KLD and Lindel perform identically, given that they are based on the same model, so the comparison is then made with FORECasT, the next best of the non-X-CRISP models.

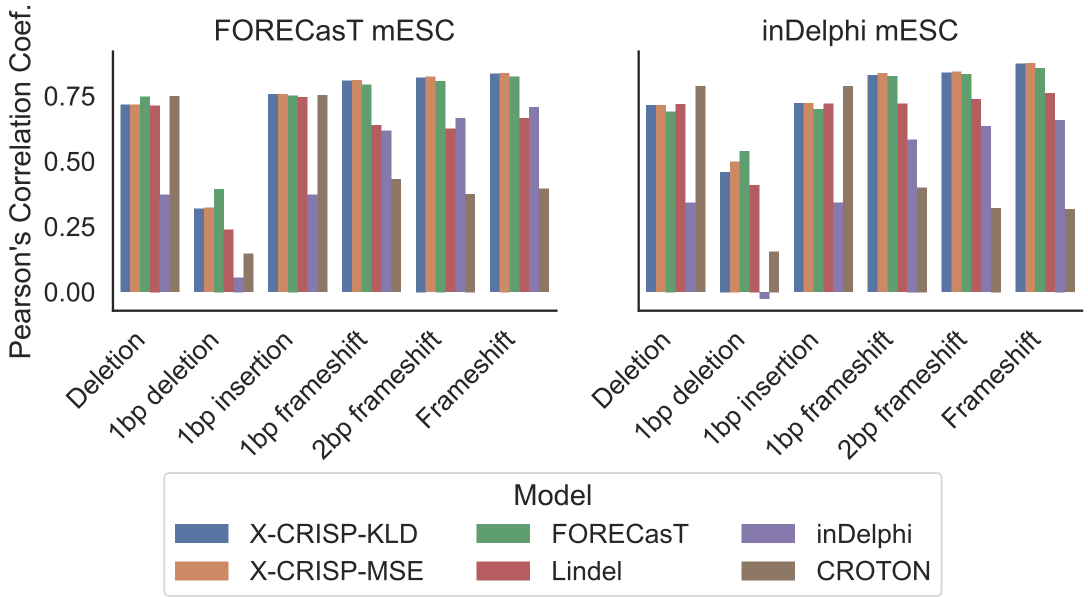

Figure S23: Broader repair outcome prediction performance. Pearson's correlation coefficient for six outcomes: deletion, 1bp insertion, 1bp deletion, 1bp frameshift, 2bp frameshift, and frameshift frequency prediction. Models trained on FORECasT WT mESC and tested on 3954 FORECasT/1961 inDelphi WT mESC target sites.

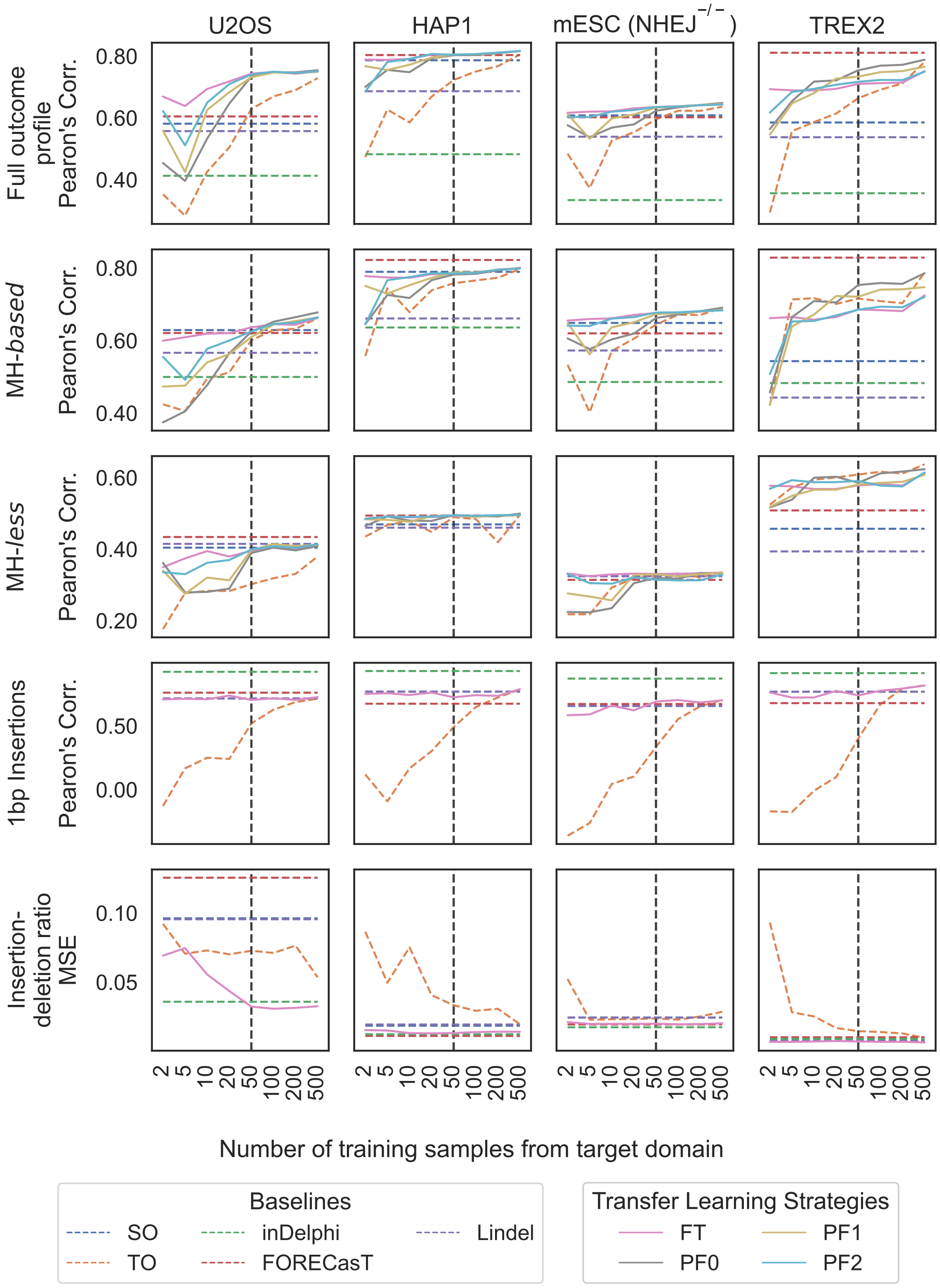

Figure S24: X-CRISP model adaptation to new domains or cell lines using transfer learning (TL). Prediction performance of baseline models and TL strategies, as the average Pearson's correlation or the MSE between predicted and observed frequencies per model and number of training samples. Baseline models: TO, X-CRISP trained on target only; SO, X-CRISP trained on source only; FORECasT, Lindel, inDelphi. Transfer learning: FT, pre-trained on source and fine-tuned for target; PF0-2, pre-trained on source and retrained + fine-tuned on target using 0-2 frozen hidden layers. (Top to bottom) Prediction models for full repair profile, MH-*based* deletions, MH-*less* deletions, insertions, and deletion-insertion ratios. Note: horizontal axis is not to scale; and the SO baseline does not use any samples from the target domain, so its performance remains constant along the horizontal axis.

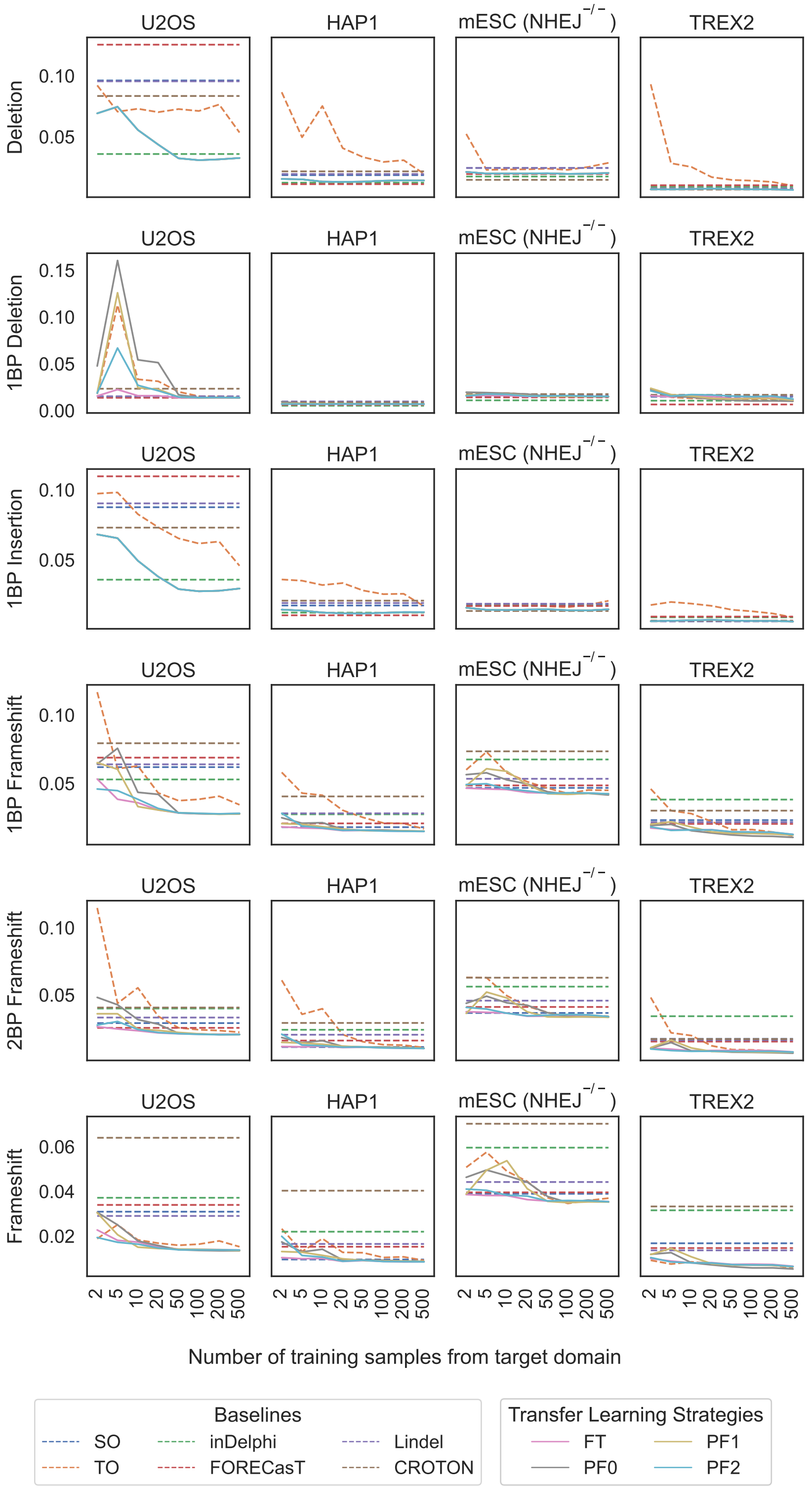

Figure S25: X-CRISP model adaptation via Transfer Learning on aggregate tasks evaluated via MSE. Prediction performance of baseline models and TL strategies, as the MSE between predicted and observed frequencies per model and number of training samples. Baseline models: TO, X-CRISP trained on target only; SO, X-CRISP trained on source only; FORECasT, Lindel, inDelphi, CROTON. Transfer learning: FT, pre-trained on source and fine-tuned for target; PF0-2, pre-trained on source and retrained + fine-tuned on target using 0-2 frozen hidden layers. (Top to bottom) Prediction models for frequency of deletions, 1bp deletions, 1bp insertions, 1bp frameshift, 2bp frameshift, and any frameshift. Note: The horizontal axis is not to scale, and the SO, FORECasT, inDelphi, Lindel, and CROTON baselines do not use any samples from the target domain, so its performance remains constant along the horizontal axis.

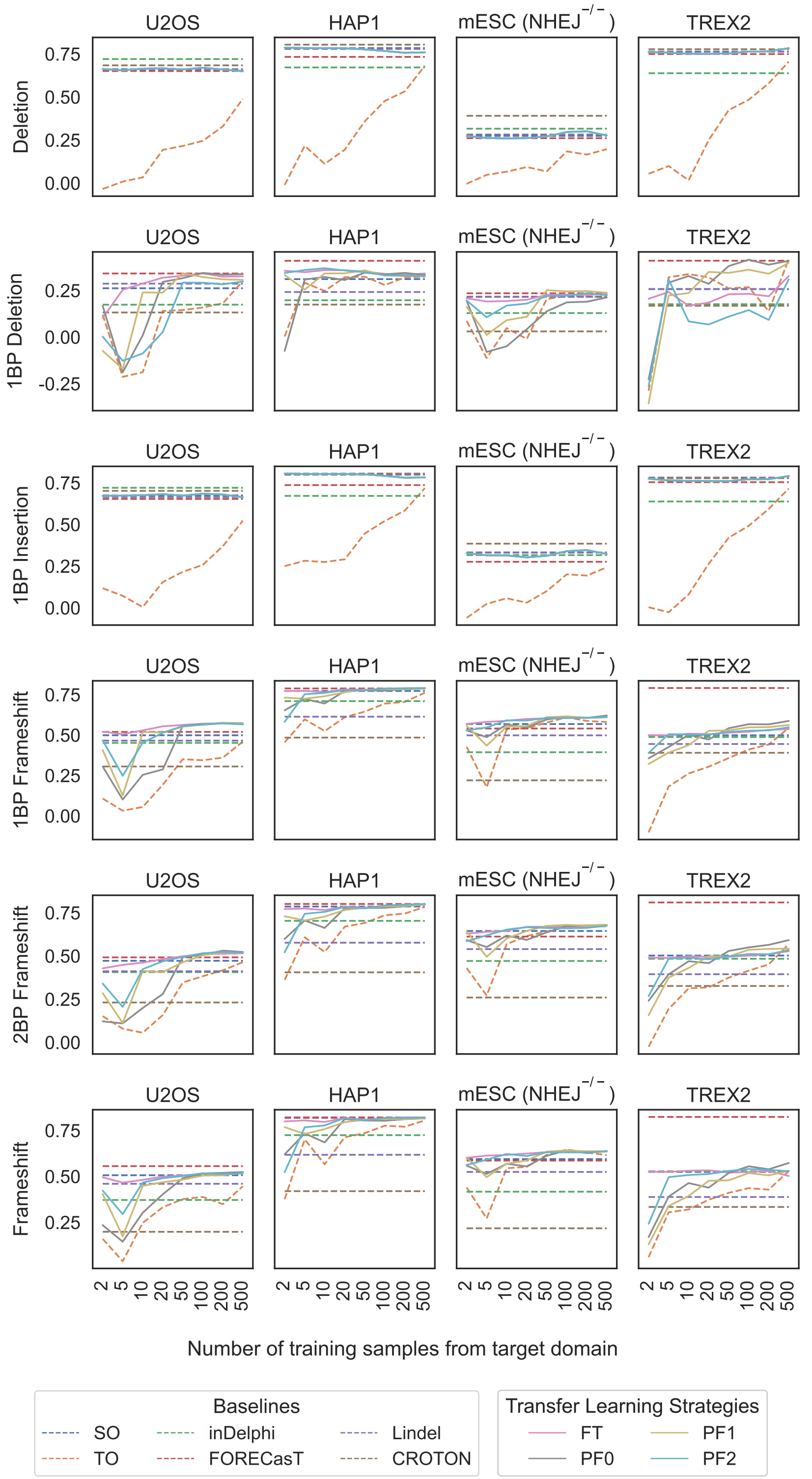

Figure S26: X-CRISP model adaptation via Transfer Learning on aggregate tasks evaluated via Pearson's correlation. Prediction performance of baseline models and TL strategies, as the Pearson's correlation between predicted and observed frequencies per model and number of training samples. Baseline models: TO, X-CRISP trained on target only; SO, X-CRISP trained on source only; FORECasT, Lindel, inDelphi, CROTON. Transfer learning: FT, pre-trained on source and fine-tuned for target; PF0-2, pre-trained on source and retrained + fine-tuned on target using 0-2 frozen hidden layers. (Top to bottom) Prediction models for frequency of deletions, 1bp deletions, 1bp insertions, 1bp frameshift, 2bp frameshift, and any frameshift. Note: The horizontal axis is not to scale, and the SO, FORECasT, inDelphi, Lindel, and CROTON baselines do not use any samples from the target domain, so its performance remains constant along the horizontal axis.

### References

[1] F. Allen, L. Crepaldi, C. Alsinet, A. J. Strong, V. Kleshchevnikov, P. De Angeli, P. Páleníková, A. Khodak, V. Kiselev, M. Kosicki, et al. Predicting the mutations generated by repair of cas9-induced double-strand breaks. *Nature biotechnology*, 37(1):64–72, 2019.

[2] W. Chen, A. McKenna, J. Schreiber, M. Haeussler, Y. Yin, V. Agarwal, W. S. Noble, and J. Shendure. Massively parallel profiling and predictive modeling of the outcomes of CRISPR/Cas9-mediated double-strand break repair. *Nucleic Acids Research*, 47(15):7989–8003, 2019.

[3] V. R. Li, Z. Zhang, and O. G. Troyanskaya. CROTON: an automated and variant-aware deep learning framework for predicting CRISPR/Cas9 editing outcomes. *Bioinformatics*, 37(Supplement_1):i342–i348, 2021.

[4] M. W. Shen, M. Arbab, J. Y. Hsu, D. Worstell, S. J. Culbertson, O. Krabbe, C. A. Cassa, D. R. Liu, D. K. Gifford, and R. I. Sherwood. Predictable and precise template-free crispr editing of pathogenic variants. *Nature*, 563(7733):646–651, 2018.
